# Chromatin is a long-range force generator that regulates plasma membrane tension and cell integrity independently of gene expression

**DOI:** 10.1101/2025.10.02.680155

**Authors:** Aidan T. Cabral, Manasi Sawant, Minwoo Kang, Xie Liangqi, Hawa Racine Thiam

**Affiliations:** Department of Bioengineering, Stanford University, California 94305; Lerner Research Institute, Cleveland Clinic, Ohio, USA; Department of Microbiology and Immunology; Sarafan ChEM-H Institute, Stanford University, California 94305

**Keywords:** Chromatin, NETosis, plasma membrane, microscopy

## Abstract

Primarily studied for its role in gene expression, chromatin organization is emerging as an important regulator of nuclear mechanics. Although the nucleus is in mechanical equilibrium with the cell, we do not know whether and how chromatin reorganization actively regulates the mechanical properties and downstream behaviors of cells. Here, we tested the hypothesis that as a dynamic crosslinked polymer, chromatin directly impacts cell mechanics independently of transcription by studying NETosis: a transcription-independent process where chromatin decompacts and the plasma membrane (PM) ruptures. Using high resolution microscopy and ATAC-see, we found that chromatin accessibility progressively increases during NETosis suggesting that chromatin binding proteins (CBPs) dissociate from chromatin during NETosis. To determine the identity and dynamics of these dissociated CBPs, we used fluorescent recovery after photobleaching to measure the mobility and localization of the linker histone H1, the nucleosomal histone H3 and the heterochromatin binding protein HP1α. We found that the mobile fraction of nuclear H1 increases during NETosis while fractions of HP1α and H3 diffuse outside of the nucleus suggesting that they become cytosolic osmolytes and potentially alter the mechanical state of cells. Consistently, we found that plasma membrane tension and cell volume increase as chromatin decompacts during NETosis. In non-NETing U2OS cells, we found that inducing chromatin decompaction increases plasma membrane tension, independently of the cytoskeleton, indicating a causal relationship between chromatin organization, cell volume and plasma membrane tension. Our findings reveal a novel non-genetic role of chromatin in cellular biophysics: regulating cell volume, PM tension, and thus, overall cell mechanics. Considering the critical role of cell mechanics in biological processes such as cell migration, proliferation and pathogen killing, our work broadens our understanding of how chromatin regulates cell physiology and pathology.

**Significance Statement:** Chromatin organizes our DNA inside the nucleus and is important for gene expression. However, chromatin is also a polymer which can passively regulate the rigidity of the nucleus, but whether and how chromatin can actively regulate the mechanical properties of the whole cell remains unknown. Here, we leverage the immune process of NETosis to show that the organization of chromatin inside the nucleus actively regulates the volume and tension of cells. Our work establishes chromatin as a long-range force generator in cells, broadening our understanding of the roles of this crucial polymer network in cells and opening the door to new strategies for controlling the mechanical properties of cells as needed by their physiology.

## Introduction

The mechanical properties of cells (stiffness, viscoelasticity, tension) actively regulate crucial biological processes including cell migration, proliferation, differentiation and pathogen killing by immune cells (1–6). These properties allow cells to sense, respond to and resist physical forces omnipresent in *in vivo* environments (6). For instance, plasma membrane tension – defined by in-line tension of the lipid bilayer and the adhesion strength between the cortex and the lipid bilayer (7)– provides mechanical resistance to the mitotic spindle in dividing cells, allowing proper chromosome segregation and successful cell division (8). Plasma membrane tension is also a long-range mechanical integrator which allows cells to cluster receptors at the surface to sense and transduce extracellular forces for optimal cell migration (9, 10). Finally, plasma membrane tension allows immune cells to spread on their targets and engage their receptors for activation and subsequent target neutralization (11, 12). The critical role of plasma membrane tension in cell physiology and disease has motivated many studies to understand the regulators of plasma membrane tension (12, 13). We thus know that plasma membrane reservoir and cytoskeletal dynamics regulates plasma membrane tension (13).

Plasma membrane reservoir allow mammalian cells to buffer tension (14). On the other hand, the cytoskeleton applies mechanical forces and regulates the composition of the plasma membrane – which impacts in-line tension - by directly binding certain lipids (15) or proteins (16) and by regulating the trafficking of vesicles from and to the plasma membrane (e.g., endocytosis/exocytosis) (17, 18). However, besides the membrane reservoirs and the cytoskeleton, we have little understanding of how other cellular components or the dynamics of other polymer networks in cells regulate plasma membrane tension. This limits our understanding of how cells actively regulate plasma membrane tension to accomplish crucial functions such as pathogen killing.

The cell nucleus is a major structural element in eukaryotic cells and is physically connected to the cytoskeleton and the rest of the cell, including the plasma membrane. This physical connection is primarily mediated by the “Linker of the nucleoskeleton and cytoskeleton” (LINC) complex (19, 20) and allows forces to be transmitted between the nucleus and the rest of the cell (21) ensuring mechanical equilibrium between these compartments. For instance, extracellular forces sensed by receptors at the plasma membrane, are transduced through the cytoskeleton and the LINC complex driving mechanical and transcriptional changes in the nucleus (22). Such mechanotransduction allows cells to interpret and adapt to mechanical signals for optimized migration, proliferation, differentiation or pathogen killing by immune cells (11, 23–26). Because of the critical role of the LINC complex and the cytoskeleton in cellular mechanotransduction, the current model proposes that the cytoskeleton is required for force transmission and the overall mechanical integration between the cell and the nucleus. However, recent work challenges this notion (27). For instance, during immune cell migration in confining microenvironments, forces from actin polymerization can be transmitted to the nucleus for its deformation, independently of the LINC complex (28). Immune cell path finding during migration in complex environments is regulated by the mechanical properties of the nucleus, independently of the LINC complex (29). Furthermore, the cell, plasma membrane and nucleus drastically remodel in synchrony and in absence of the cytoskeleton during neutrophil extracellular trap release (NETosis) (30). How forces are transmitted between the nucleus and the plasma membrane independently of the cytoskeleton is an open question that studying NETosis can help us answer.

NETosis is a transcription-independent (31, 32) evolutionary conserved (33, 34) immune response during which neutrophils release chromatin to the extracellular space to form neutrophils extracellular traps (NETs) which can ensnare and kill pathogens but also propagate inflammation (35). NETosis can take two forms: suicidal NETosis that leads to neutrophil death (36, 37), and vital NETosis where neutrophil remain alive (38). Suicidal NETosis, the best studied form of NETosis, proceeds via a well conserved sequence of cellular events (30) that starts with disassembly of the actin, microtubules and vimentin cytoskeleton, before drastic remodeling of the nucleus and plasma membrane. Nuclear remodeling during NETosis is characterized by chromatin reorganization, nuclear rounding and final nuclear rupture. Plasma membrane remodeling is characterized by shedding of microvesicles and final plasma membrane rupture (30). We have limited understanding of how plasma membrane remodeling is regulated or predispose the membrane for rupture, the final critical step of NETosis required for NET release.

For instance, a hallmark of NETosis is the drastic reorganization of chromatin, the longest polymer network in eukaryotes (39). During NETosis, chromatin accessibility (40) and the area occupied by chromatin in cells (41) increases. Further, chromatin in NETs is reported to have a “beads-on-string” structure suggesting that chromatin is decompacted down to its nucleosomal level of organization (42). Histone posttranslational modifications (PTMs) namely citrullination of histone H3 at Arg2, Arg8, Arg17 and histone H4 at Arg3 (43) by Peptidylarginine deiminase 4 (PAD4) enzyme (44) and cleavage of histones H1, H2A, H2B, H3, and H4 by serine proteases such as neutrophil elastase and Proteinase 3 (PR3) (45, 46) are proposed to drive chromatin decompaction during NETosis by dissociating histones from DNA (35).

Decompacted chromatin is proposed to expand and fill the cell, exerting entropic elastic pressure on the plasma membrane that ultimately drives rupture during NETosis (41). However, the dynamics and impact of chromatin reorganization inside the nucleus during NETosis remains unclear. We do not know whether, when and which chromatin binding proteins, including histones, dissociate from chromatin. We also do not know whether chromatin reorganization, which has been shown to alter the mechanical properties of the nucleus (47–52), impacts the mechanical properties and integrity of the plasma membrane and the cell overall, especially in the absence of the cytoskeleton.

Here we leverage the transcription-independent process of NETosis to test the hypothesis that the nucleus, in particular chromatin directly regulates plasma membrane tension and the overall mechanical state of cells, in the absence of the cytoskeleton and independently of gene expression. We found that chromatin binding proteins that impart chromatin structure (H1, HP1α, H3), rapidly and progressively dissociate from chromatin as NETosis progresses.

We found that these dissociated chromatin binding proteins become more soluble in the nucleus and the cell, increasing the concentration of macromolecules in the cell. We posit that this increase in soluble macromolecules due to chromatin decompaction increases cellular osmotic pressure resulting in an increase in volume and plasma membrane tension. Consistently, we found that cell volume and plasma membrane tension increase as NETosis progresses. We showed that this chromatin decompaction induced increase in plasma membrane tension is a general biophysical mechanism that we could recapitulate in non-NETing cells. Inducing decompaction in non-immune cells increases plasma membrane tension independently of the cytoskeleton. Together, our data reveals a novel way in which chromatin organization directly regulates the mechanical state of the cell, independently of the cytoskeleton.

## Results

### Chromatin accessibility progressively increases from the early stages of NETosis

#### Chromatin homogenizes from the early stages of NETosis

To determine the dynamics of chromatin reorganization throughout NETosis, we performed high resolution spinning-disc confocal microscopy of live HL60-derived neutrophils (dHL60 cells) stained with a fluorescent DNA dye (SPY650-DNA) and stimulated to undergo NETosis using a calcium ionophore (Ionomycin; 4 µM). We previously showed and here confirmed (**Figure 1A – C, Movie S1**), that dHL60 cells stimulated with 4 µM of ionomycin undergo NETosis using the same cellular mechanism as primary human and mouse neutrophils stimulated for NETosis with Ionomycin, the bacterial derivative lipopolysaccharide (LPS) or live yeast (Candida Albicans) (30). This cellular mechanism proceeds as follows: shedding of microvesicles from the plasma membrane and onset of cell rounding, reorganization of the actin cytoskeleton leading to its depolymerization (**Figure S1, Movie S2**), reorganization of chromatin, rounding of the nucleus, rupture of the nuclear envelope which allows expansion of chromatin in the cytosol, permeabilization and rupture of the plasma membrane which allows NETosis completion via release of chromatin to the extracellular space (**Figure 1 A-C**; see Materials and Methods). Consistent with our previous work, we defined NETosis onset as the first time point when plasma membrane microvesicles are detected or cells start rounding up.

**Figure 1:**
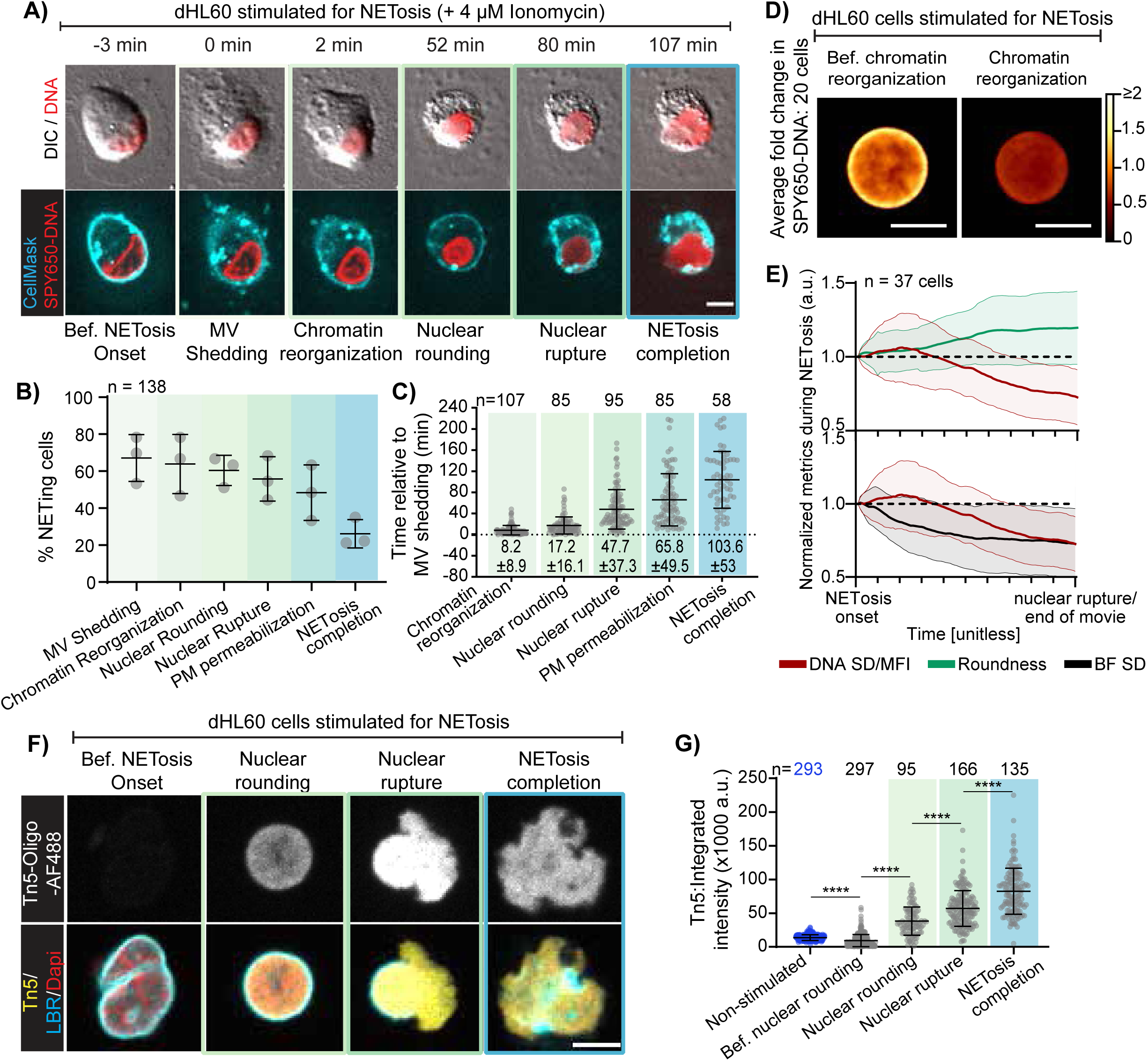
Chromatin accessibility increases progressively from the early stages of NETosis. dHL60 cells stained with SPY650-DNA were stimulated with ionomycin (4 µM,) and imaged by DIC and spinning confocal microscopy **(A-C, E-G)** or laser scanning microscope **(D)** at the cell-coverslip interface (Z= 0 μm) and 3 μm above **(A-C)** (Z =+3 μm) at 1 min intervals for 4h. **(A)** Representative montage of a dHL60 cell at different timepoints following ionomycin stimulation. Images show overlays of SPY650-DNA fluorescence (red) with DIC (grayscale, top row) and cell membrane dye fluorescence (CellMask, cyan, bottom row). Colored boxes highlight cellular events corresponding to the graphs in **(B)**, **(C)** and **(G)**. **(B)** Percentage of cells undergoing microvesicle (MV) shedding, chromatin reorganization, nuclear rounding, nuclear rupture (DNA release into the cytoplasm), plasma membrane (PM) permeabilization (loss of DIC contrast at the cell periphery), and NETosis completion (extracellular DNA release) after stimulation. n = total number of cells; each data point represents a biological replicate. **(C)** Timing of cellular events initiation relative to MV shedding. n = total number of cells, each data point indicates an individual cell. **(D)** Average paired heatmaps of SPY650-DNA in single NETing cells at early (left) and late (right) stages of chromatin reorganization. Heatmap indicates relative fold change in SPY650-DNA dye intensity. n = 20 cells. **(E)** Quantification of nuclear roundness, the standard deviation (SD) of SPY650-DNA normalized to the DNA mean fluorescence intensity (MFI) (DNA SD/MFI), and the standard deviation (SD) of the brightfield channel of the nucleus (BF SD), between NETosis onset (MV shedding or onset of cell rounding) and 1 min before nuclear rupture or end of the movie. Metrics were normalized to their initial value at NETosis onset. Bold lines: mean; thin lines: ± standard deviation; Top panel: DNA SD/MFI (red) and nuclear roundness (green); Bottom panel: DNA SD/MFI (red) and BF SD (black). n = 37 cells from 2 experimental replicates. **(F)** Representative images of TN5 Oligo-AF488 only (top) or overlayed with DAPI and the Lamin B Receptor (LBR; Bottom) in dHL60 cells at different stages of NETosis, from 3D Z stack; step size 0.3 µm and range 9 µm. **(G)** Quantification of total fluorescence intensity of Tn5-Oligo-AF488 in cells at different stages of NETosis as assessed by nuclear shape (roundness), nuclear envelope integrity (continuous or discontinuous LBR staining). n = total number of cells; Each datapoint indicates individual cell. Brackets denote the presence of 4 μM ionomycin in **(A)**, **(D)** and **(F)**. Bars in **(B)**, **(C)** and **(G)** indicate mean ± SD. Statistical test: Mann-Whitney U test (two-tailed, non-parametric). Analysis was performed using GraphPad Prism v10. Scale bar in **(A)**, (**D)** and **(G)**: 5 μm.

Nuclear and plasma membrane rupture was defined as the first timepoint when we detect DNA exit from the nucleus or cell, respectively. Analysis of the percentage and timing of NETosis progression in control dHL60 cells and in cells treated with the transcription inhibitor, actinomycin D (1mg/mL), or the translation inhibitor, cycloheximide (100 nM), showed that transcription- or translation-inhibited cells undergo NETosis at similar rates (**Figure S2A, C**) and following the same sequence of cellular events (**Figure S2B, D**) than control dHL60 cells (**Figure S2**). This data confirms previous reports (53, 54) that NETosis initiation and completion does not require de novo gene expression or protein synthesis. This suggests that the mechanism and impact of chromatin reorganization in NETosis are independent of the transcription roles of chromatin.

To start probing the dynamics of chromatin reorganization during NETosis, we assessed the distribution of nuclear signal between NETosis onset and nuclear rupture. First, we quantified changes in the brightfield image of the nucleus as well as the distribution of a DNA dye (SPY650-DNA) by measuring the standard deviation (SD) of the brightfield channel (BF SD) and SD of DNA divided by its mean fluorescence intensity (DNA SD/MFI) from time-lapse spinning disc confocal movies of dHL60 cells undergoing NETosis (**Figure 1E, S3A-E, H; Movie S3;** see Materials and Methods). This showed that both BF SD and DNA SD/MFI decrease by 27% (**Figure 1E & S3H**) between NETosis onset and nuclear rupture suggesting that nuclear contents, including chromatin, homogenizes in the nucleus prior to nuclear rupture. To determine how DNA/chromatin homogenization relates to changes in nuclear shape, we calculated nuclear roundness between NETosis onset and nuclear rupture (**Figure S3 F-H**) and analyzed its temporal evolution relative to DNA SD/MFI (**Figure 1E-top**). This revealed that the increase in nuclear roundness (**Figure S3F-H**) precedes the decrease in DNA SD/MFI (**Figure 1E-top**) indicating that DNA homogenizes after the nucleus rounds up during NETosis. The lack of temporal correlation between DNA homogenization and nuclear rounding further suggests that DNA homogenization is not a mere result of the change in nuclear shape. DNA homogenization could be due to the whole chromatin decompacting into loosely packed euchromatin or compacting into densely packed heterochromatin. To start probing the packing state of chromatin, we measured changes in mean fluorescence intensity of nuclear SPY650- DNA (DNA MFI) between NETosis onset and nuclear rupture (**Figure S3 E, H**). This showed that DNA MFI decreases by ∼15%, suggesting that chromatin packing decreases (**Figure S3H**). To map the spatial distribution of DNA homogenization, we projected the DNA MFI of 20 nuclei of NETing cells with visually round nuclei before and after DNA/chromatin homogenization (**Figure 1D**, see Materials and Methods). Analysis of this population average DNA MFI map revealed that DNA MFI at the nuclear periphery decreases by >2 fold, suggesting that DNA homogenization is initiated at the nuclear periphery. Since DNA is the founding unit of chromatin, this data suggests that chromatin spatially and structurally reorganizes to homogenize inside the nucleus from the early stages of NETosis.

#### Chromatin accessibility increases as chromatin homogenizes inside the nucleus at the early stages of NETosis

To assess whether early chromatin homogenization translates into changes in molecular structure of chromatin during NETosis, we used assay of transposase-accessible chromatin with visualization (ATAC-see (40) to measure chromatin accessibility as cells progress towards NETosis (**Figure 1F-G, S3I;** see Materials and Methods). Non-stimulated and NETosis-stimulated dHL60 cells (60 min of stimulation) were fixed and stained with the transposon enzyme Tn5 conjugated with a DNA oligo tagged with the fluorescent dye Alexa Fluor 488 (will be concisely referred to as Tn5). Tn5 labelled cells were then stained with DNA dyes (SPY650- DNA and DAPI) and antibodies against a nuclear envelope protein (Lamin B Receptor, LBR).

Stained cells were imaged at high spatial resolution using a spinning disc confocal microscope. We acquired 3D stacks to capture the entire cell and nucleus. From the 3D Z-stack images, cells were classified in different stages of NETosis depending on nuclear shape (i.e., before or after nuclear rounding), the integrity of the nuclear envelope (i.e., before or after nuclear rupture) and the integrity of the cell (i.e., before or after plasma membrane rupture / NETs release resulting in NETosis completion) (**Figure 1A,F**). Quantification of the total fluorescence intensity of Tn5 on DNA for each single cell (see Materials and Methods) showed that the Tn5 intensity progressively increases as cells progress towards NETosis (**Figure 1 G**). Notably, Tn5 intensity significantly decreased in the early stages of NETosis (**Figure 1G**: comparing “Non-stimulated” and “Before nuclear rounding” groups) before increasing ∼4-fold in cells at the nuclear rounding stage of NETosis. Tn5 intensity further increases after nuclear rupture (∼6-fold compared to “Before nuclear rounding”) and NETs release (∼9-fold compared to “Before nuclear rounding”) (**Figure 1F, G**). To determine how increase in nuclear Tn5 relates to DNA/chromatin homogenization, we performed a correlation analysis between Tn5 intensity and DNA SD/MFI for NETing cells with round and intact nucleus (nuclear rounding stage of NETosis) (**Figure S3I**, see Materials and Methods). This revealed a significant negative correlation between these two parameters indicating that chromatin accessibility increases as chromatin homogenizes inside the nucleus during NETosis. This data shows that chromatin accessibility dynamically changes during NETosis starting from the early stages of the process. The progressive increase in chromatin accessibility before nuclear rupture (**Figure 1F, G**) suggests that “old” chromatin binding proteins progressively dissociate from chromatin prior to nuclear rupture during NETosis.

#### Chromatin structural proteins sequentially dissociate from chromatin and change their mobility during NETosis

We next sought to identify the chromatin binding proteins that dissociate from chromatin during NETosis. The increase in chromatin accessibility and homogeneity suggests a transition from heterochromatin to euchromatin structures. Thus, we sought to measure the dynamics of chromatin binding proteins known to regulate the 3D structure of chromatin and mediate heterochromatin formation. Specifically, we probed the mobility of H3 - a nucleosomal histone proposed to dissociate from chromatin following its citrullination by PAD4 (55); H1 - the linker histone which binding on chromatin allowing the formation of higher chromatin orders (56); and HP1α - a major organizer of heterochromatin (57) (**Figure 2A**).

**Figure 2:**
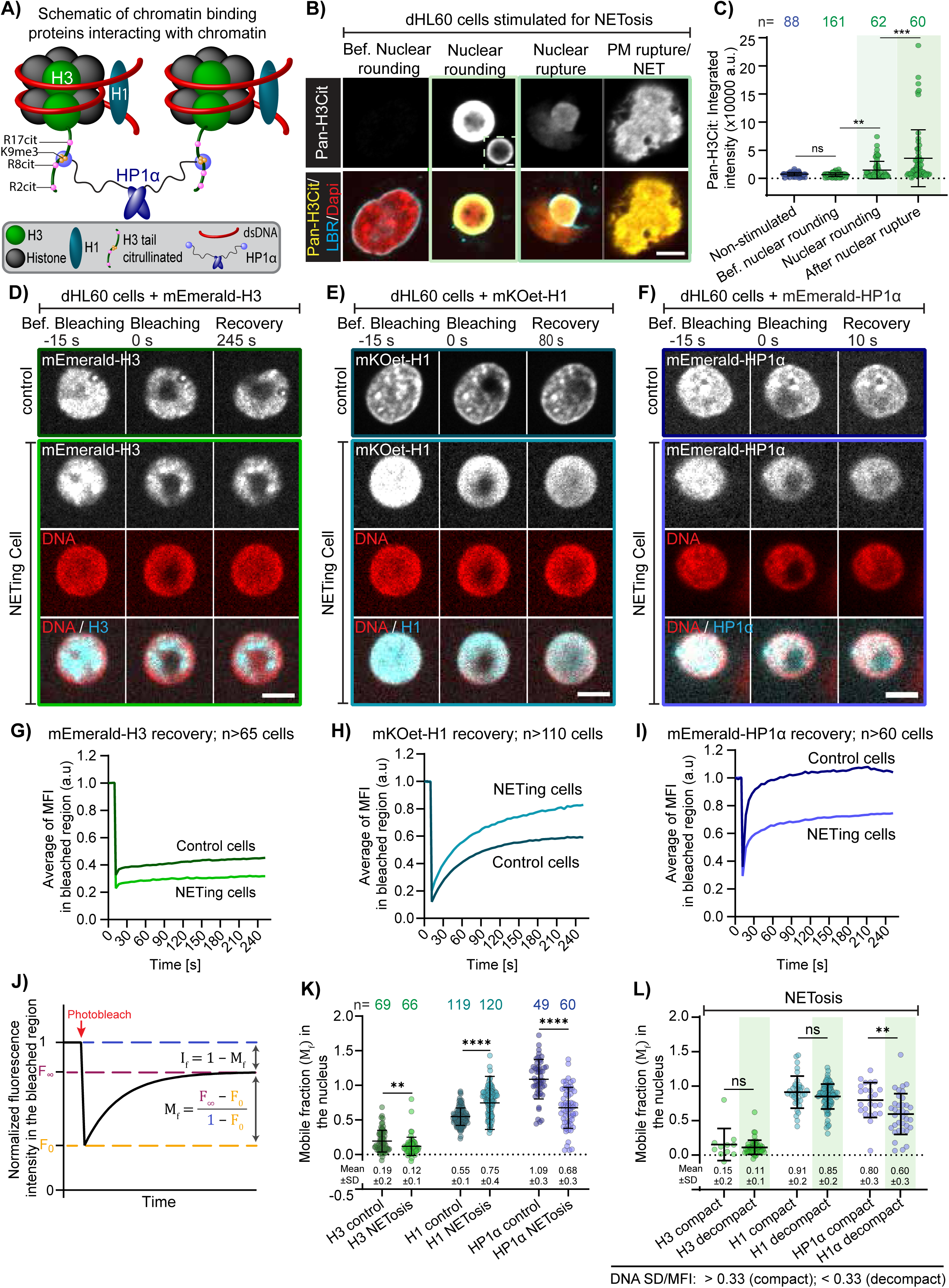
H1 dissociates from chromatin while nuclear H3 and HP1α become less mobile during NETosis. **(A)** Schematic representation of chromatin-binding proteins of interest (H3, H1, HP1α) interacting with nucleosomal chromatin. Nucleosomes consist of histone H3 (green) and other core histones H2A, H2B, H4 (gray) wrapped by double-stranded DNA (red). Linker histone H1 (teal) binds at the DNA entry/exit site of nucleosomes. The histone H3 N-terminal tail carries multiple post-translational modifications, including citrullination of arginines (R2cit, R8cit, R17cit; pink) and trimethylation of lysine 9 (K9me3; yellow). HP1α (purple) interacts with H3K9me3 to stabilize chromatin structure**. (B)** Representative immunostaining images of Pan-H3Cit only (top) or overlayed with DAPI and the lamin B receptor (LBR; Bottom) in dHL60 cells at different stages of NETosis, from 3D Z stack; step size 0.3 µm and range 9 µm. Inset in “nuclear rounding” represent a lower contrast version of the same image. **(C)** Quantification of the integrated intensity of DNA-associated Pan-H3Cit of cells at different stages of NETosis as assessed by nuclear shape (roundness), nuclear envelope integrity (continuous or discontinuous LBR staining). Each datapoint represents a cell. **(D-F)** Representative montages of FRAP experiments for control and NETing dHL60 cells expressing mEmerald-H3 **(D)**, mKOet- H1 **(E)**, and mEmerald-HP1α **(F)**. DNA is labeled with SPY650-DNA (red), and fluorescently tagged proteins are shown in cyan/white. **(G–I)** Recovery curves showing the average mean fluorescence intensity (MFI) of the bleached nuclear region (∼3 μm²). Curves were corrected for background and normalized to the fluorescence intensity in the unbleached, reference region. Darker colors represent control cells, lighter shades represent NETing cells. **(J)** Schematic of mobile fraction calculation (see Materials and Methods). **(K)** Quantification of the mobile fraction of mEmerald-H3 (green), mKOet-H1 (teal), and mEmerald-HP1α (purple) in the nucleus of control and NETing cells; mean ± SD shown below. n: total number of cells; Each datapoint indicates a cell. **(L)** Mobile fraction of mEmerald-H3 (green), mKOet-H1 (teal), and mEmerald- HP1α (purple) in the nucleus of NETing cells shown in **(K)** stratified into compact versus decompacted chromatin (light green shaded rectangle) based on K-means clustering of DNA SD/MFI (compact > 0.33; decompact < 0.33); mean ± SD shown below. Brackets denote the presence of 4 μM ionomycin in **(B)**, **(D)**, **(E)**, **(F)**, and **(L)**. Bars in **(C)**, **(K)** and **(L)** indicate mean ± SD; Statistical test: Mann–Whitney U test (two-tailed, non-parametric); ns: not significant; **: p-value < 0.01 ****: p-value < 0.0001. Scale bar: 5 μm. Analysis performed with GraphPad Prism v10.

#### Histone H3 is citrullinated in early stages of NETosis but does not increase in mobility inside the nucleus

To determine whether and when H3 dissociates from chromatin during NETosis, we first investigated the timing of histone H3 citrullination, a modification proposed to precede and drive H3 dissociation from chromatin (43). To investigate the timing of H3 citrullination relative to chromatin accessibility changes during NETosis, we fixed and stained non-stimulated and NETosis-stimulated dHL60 cells (60 min of stimulation, see Materials and Methods). Global histone H3 citrullination was assessed using a pan-H3Cit antibody that recognizes citrullinated arginine residues at positions Arg2, Arg8, and Arg17. Cells were also stained with a DNA dye (DAPI) to assess chromatin homogenization and immunostained for a nuclear envelope protein (LBR) to assess nuclear shape, nuclear envelope integrity (ruptured or not) and NETosis progression. High resolution 3D Z-stack of stained cells were acquired using a spinning disc confocal microscope. Visualization (**Figure 2B**) and analysis (**Figure 2C, S4,** see Materials and Methods) of the fluorescence intensity of pan-H3Cit in non-stimulated versus NETosis-stimulated cells revealed that H3 citrullination is non-detectable in non-stimulated cells (**Figure 2C and S4A**). We first detected citrullinated H3 at the nuclear periphery of NETing cells with round and intact nuclei (**Figure 2 B,C**; compare “Bef. nuclear rounding” and “Nuclear rounding”), indicating that H3 citrullination is an early stage of chromatin reorganization and occurs before the rupture of the nuclear envelope during NETosis. Quantitative analysis of NETing cells showed a ∼2 and ∼4-fold increase in mean (**Figure S4A**) and total (**Figure 2C**) pan-H3Cit fluorescence intensity between NETosis onset (labelled as “Bef. nuclear rounding”) and nuclear rounding (**Figure 2B and S4A**). Pan-H3Cit mean and total fluorescent intensity further increases by ∼5.5-fold after nuclear rupture, relative to NETosis onset (**Figure 2 B, C, S4A**). To determine whether H3 citrullination coincides with chromatin homogenization, we performed correlation analysis between pan-H3cit intensity and DNA SD/MFI (**Figure S4**). This revealed a significant negative correlation between these two parameters (**Figure S4B, C**), indicating that pan-H3Cit intensity increases as chromatin homogenizes inside the nucleus. This data shows that H3 citrullination starts at the nuclear periphery of NETing cells with intact nuclei and correlates with chromatin homogenization and the increase in chromatin accessibility.

We next sought to determine whether H3 dissociates from chromatin during NETosis by measuring the mobile fraction of fluorescently tagged H3 (mEmerald-H3) using fluorescence recovery after photobleaching (FRAP) (**Movie S4**; see Materials and Methods). dHL60 cells were transfected with mEmerald-H3 and left to recover for over 10 hours to ensure incorporation of mEmerald-H3 into chromatin. Cells were stained with SPY650-DNA to visualize the nucleus then FRAPed using a spinning disc confocal microscope equipped with a FRAP module. A 3 µm^2^ region of interest (ROI) at the center of the nucleus of non-stimulated or NETosis-stimulated cells was bleached using a 405 nm laser (dark region of the nucleus in **Figure 2D – F**). Since H3 citrullination and increase in chromatin accessibility was first measured in NETing cells with round nuclei, we focused on measuring the mobility of H3 in NETing cells with round nuclei (Roundness = 0.95 +/-0.04; **Figure S5A**). From the reference and background corrected fluorescence recovery curves of mEmerald-H3 (**Figure S5B, C**), we calculated the mobile fraction of nuclear H3 (see Material and Methods; **Figure 2J**). Analysis of the recovery curves and mobile fraction data shows that ∼80% of mEmerald-H3 is immobile in non-stimulated cells (**Figure 2 G, K**), indicating that mEmerald-H3 is well incorporated into chromatin. Comparison of the recovery curves and mobile fractions of mEmerald-H3 in non-stimulated versus NETosis- stimulated cells shows that the mobile fraction of H3, slightly but significantly, decreases from 0.19+/-0.2 in non-stimulated cells to 0.12+/-0.1 in NETing cells (**Figure 2K**). This decrease in the mobile fraction of H3 in cells at the nuclear rounding stage of NETosis when H3 citrullination (**Figure 2B, C**), and chromatin accessibility (**Figure 1F, G**), increased suggests that although citrullinated, H3 might remain associated with decompacted chromatin during NETosis.

Consistently, correlation analysis between the mobile fraction of nuclear mEmerald-H3 and DNA SD/MFI, at the single cell level revealed no correlation between these two parameters (**Figure S6A**), suggesting that the H3 mobility is independent of chromatin homogenization during NETosis. Furthermore, comparing the mobile fraction of mEmerald-H3 between NETing cells with homogenized and non-homogenized chromatin (DNA SD/MFI < 0.33 versus > 0.33, as determined by K-means clustering, **Figure S6D, E,** see Materials and Methods) showed no significant difference in the mobile fraction of mEmerald-H3 as function of chromatin homogenization (**Figure 2L**).

Together, this data indicate that H3 is citrullinated early during NETosis but H3 mobility remains low inside the nucleus of NETing cells with round and intact nucleus, independently of chromatin homogenization and the increase in chromatin accessibility during NETosis. Although this data contrasts with the prevailing model that H3 citrullination promotes its dissociation from chromatin, it aligns with prior studies reporting that nucleosomes remain present on extruded chromatin during NETosis (38).Together, this data indicate that H3 is citrullinated early during NETosis, but shows that H3 mobility remains low inside the nucleus of NETing cells with round and intact nucleus, independently of chromatin homogenization and the increase in chromatin accessibility during NETosis.

#### Chromatin structural proteins that mediate higher order chromatin structure change in mobility during NETosis

We then sought to determine if the linker histone H1 which organizes nucleosomes (58) dissociates from chromatin during NETosis. To probe the interaction between H1 and chromatin during NETosis, we used FRAP to analyze the mobile fraction of fluorescently-tagged H1 (mKOet-H1, **Figure 2E; Movie S5**; see Materials and Methods). dHL60 cells were co-transfected with mKOet-H1 and mEmerald-H3 to measure and compare the relative changes in recovery curves (**Figure S5 B-E**) and mobile fractions (**Figure 2K**) of H1 and H3 histones at single cell resolution. Comparison of the recovery curves and mobile fractions of mEmerald-H3 and mKOet-H1 in non-stimulated cells showed that H1 is more mobile than H3 (mobile fraction of 0.55+/-0.1 for H1 versus 0.19+/-0.2 for H3) (**Figure 2G, H, K**), consistent with the linker histone H1 having a higher mobile pool or lower affinity for chromatin than nucleosomal histones (59). The mobile fraction of mKOet-H1 further increases in NETing cells going from 0.55+/-0.1 in non-stimulated cells to 0.75+/-0.4 in NETing cells with round and intact nuclei (**Figure 2H, K**), indicating that H1 dissociates from chromatin during NETosis. Notably, comparing the mobile fraction of cells with homogenized and non-homogenized chromatin (DNA SD/MFI < 0.03 versus > 0.33, as determined by K-mean clustering, **Figure S6D, E,** see Materials and Methods), revealed no significant differences (**Figure 2L**). Furthermore, correlation analysis showed no correlation between the mobile fraction of mKOet-H1 and chromatin homogenization (**Figure S6B**). This data shows that the mobile fraction of H1 increases in early stages of NETosis and remains high, independently of chromatin homogenization and the increase in chromatin accessibility during NETosis.

We next sought to determine if HP1α, a chromatin binding protein that mediates higher order chromatin organization such as heterochromatin (56), dissociates from chromatin during NETosis. We used FRAP to measure the mobile fraction and diffusion behavior of fluorescently-tagged HP1α (mEmerald-HP1α) in non-stimulated and NETosis-stimulated dHL60 cells (**Figure 2F; Movie S6**; see Materials and Methods). dHL60 cells were co-transfected with mKOet-H1 and mEmerald-HP1α to compare the relative changes in mobility of H1 and HP1α at single cell resolution. Cells were photobleached using a 488 laser to minimize HP1α recruitment to UV-exposed DNA (60). We found that the mobile fraction of HP1α is higher than that of H1 in non-stimulated cells (1.09+/- 0.3 for HP1α versus 0.55+/-0.1 for H1) (**Figure 2K**), indicating that HP1α has a larger soluble pool or less affinity for chromatin than H1. Comparing the recovery curves (**Figure S5F, G**) and the mobile fractions of HP1α in non-stimulated cells versus NETing cells, showed that the mobile fraction of HP1α decreases by ∼40% during NETosis (**Figure 2F, I, K**). To determine whether the decrease in the mobile fraction of HP1α coincides with chromatin homogenization during NETosis, we performed single cell correlation analysis between the mobile fraction of HP1α and DNA SD/MFI. This revealed a significant negative correlation between these two parameters (**Figure S6C**), indicating that the mobile fraction of HP1α decreases as DNA homogenizes inside the nucleus. Furthermore, comparing the mobile fraction of HP1α between NETing cells revealed a ∼1.33-fold decrease in HP1α mobile faction between cells with homogenized (DNA SD/MFI < 0.33) and non-homogenized chromatin (DNA SD/MFI> 0.33) (**Figure 2L**). This data shows that the mobile fraction of nuclear HP1α decreases as chromatin decompacts during NETosis.

Together, this data indicates that the interaction between chromatin and chromatin structural proteins changes during NETosis. Specifically, the linker histone H1 which connects nucleosomes and is required for higher order chromatin organization, is more mobile inside the nucleus while the mobility of nucleosomal histone H3 or proteins that mediate higher order chromatin structure (HP1α) decreases in the nucleus of NETing cells. While the increase in H1 mobility indicates that H1 dissociates from chromatin during NETosis, the decrease in H3 or HP1α mobility could indicate their increased association with chromatin or exit of their mobile/dissociated pools from the nucleus. We set out to untangle these two possibilities.

### Dissociated chromatin structural proteins H3 and HP1α translocate from the nucleus to the cytosol

To determine if the decrease in mobile fractions of H3 and HP1α is due to soluble/dissociated H3 and HP1α exiting the nucleus in NETing cells, we analyzed the distribution and localization of fluorescently-tagged H3 and HP1α as well as endogenous HP1α during NETosis (**Figure 3, S7, S8**). dHL60 cells were transfected with mEmerald-H3 or mEmerald-HP1α and stained with SPY650-DNA and CellMask to visualize the nucleus and cell membrane, respectively. Live cells were imaged for 4 hours at high resolution by spinning disc confocal microscopy. Cells were stimulated for NETosis 3 to 6 minutes after the onset of live imaging to capture their transition into NETosis. We acquired confocal images at the interface between the cell and the coverslip (Z = 0) to visualize the dynamics of the cell membrane and at 3 µm above the coverslip (Z = 3 µm) to visualize the dynamics of the nucleus. We used the confocal image at Z = 0 to assess the onset of microvesicle shedding or cell rounding which represents our earliest detection of NETosis initiation (**Figure 1** and (30)). We used the confocal image at Z = 3 µm to assess changes in nuclear shape and integrity (e.g. nuclear rounding, DNA release into the cytosol) and quantify the distribution and localization of nuclear proteins.

**Figure 3:**
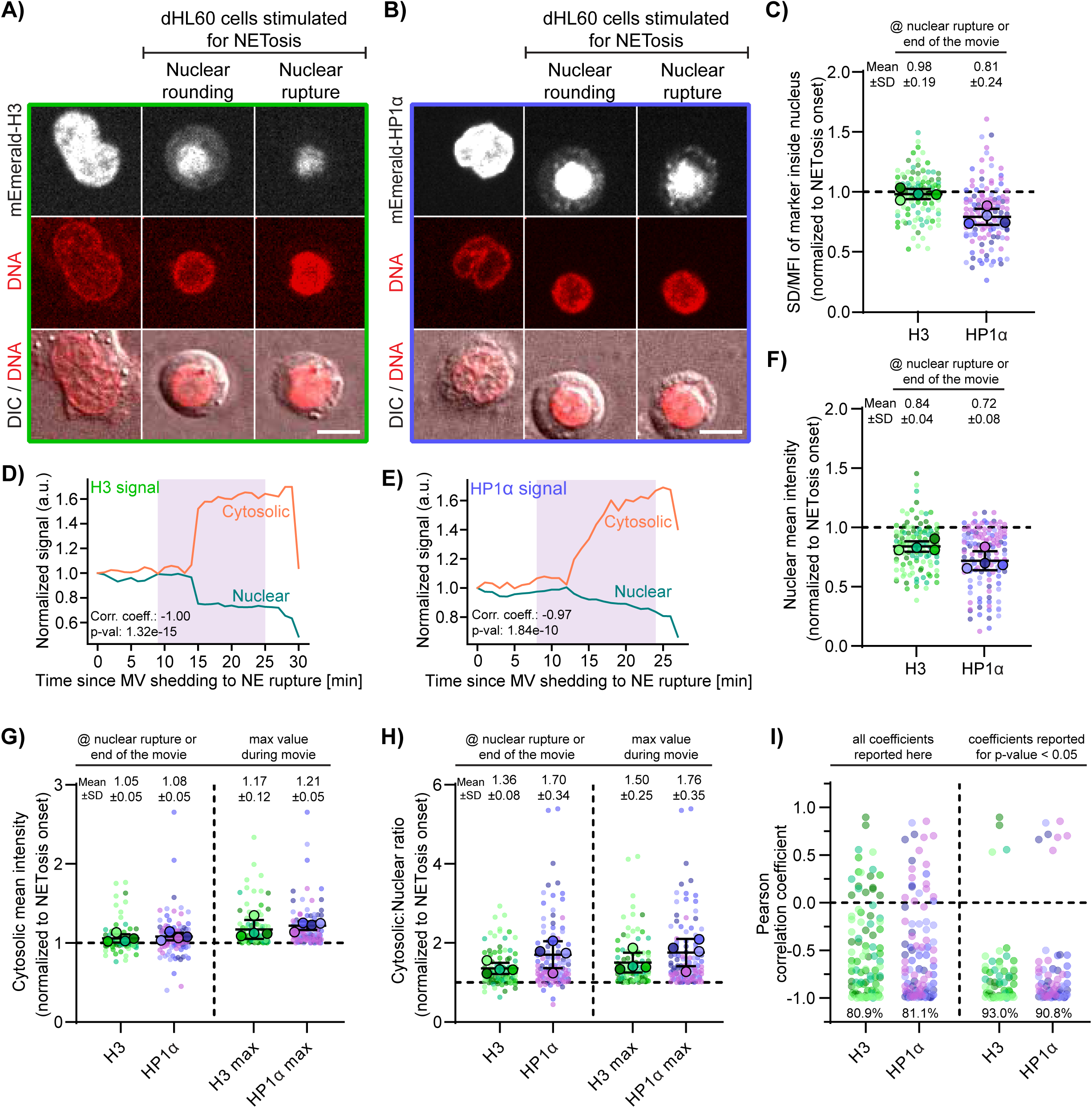
Dissociated chromatin structural proteins H3 and HP1α translocate to the cytosol (A-B) Representative montages of dHL60 cells expressing mEmerald-H3 **(A)** or mEmerald- HP1α **(B)** at defined NETosis stages. Images show the fluorescent protein (greyscale; top), DNA (red; middle), and a DIC/DNA overlay (bottom). Brackets above images denote the presence of ionomycin (4 μM). Scale bar: 5 μm. **(C)** Quantification of the standard deviation (SD) over mean fluorescence intensity (MFI) of mEmerald-H3 (green) and mEmerald-HP1α (purple) of single NETing cells at nuclear rupture or end of the movie (whichever occurred first), relative to their respective values at NETosis onset (indicated by MV shedding or onset of cell rounding). **(D)** Quantification of mean fluorescence intensity of nuclear (blue) and cytosolic (orange) mEmerald-H3 of NETing dHL60 cell in **(A)**. **(E)** Quantification of mean fluorescence intensity of nuclear (blue) and cytosolic (orange) mEmerald-HP1α of NETing dHL60 cell in **(B)**. **(D** and **E)**, nuclear and cytosolic intensities were normalized to their values at NETosis onset (MV shedding, T=0 min). Purple boxes indicate a 16-minute time window used for Pearson correlation analysis. Corresponding correlation coefficients (corr. coeff.) and p-values are reported in each plot. Note: decreased cytosolic signal reflects plasma membrane permeabilization. **(F-H)** Quantification of the nuclear **(F)** and cytosolic **(G)** mean fluorescence intensity as well as cytosolic to nuclear ratio **(H)** of mEmerald-H3 (green) and mEmerald-HP1α (purple) of single NETing cells at time points indicated above graphs relative to their respective values at NETosis onset (indicated by MV shedding or onset of cell rounding). Data in **(C)**, **(F)**, **(G)** and **(H)** shown as Superplots (84) where each small dot represents a cell; each large dot represents the average metric in a population of cells within the same biological replicate depicted with matching shades of green (mEmerald-H3) or purple (mEmerald-HP1α); Bars represent Mean ± SD of the average metric across 4 biological replicates, shown above graphs. Max: the maximum value of the corresponding metric. **(I)** Pearson correlations between nuclear and cytosolic intensities of mEmerald-H3 (green) and mEmerald-HP1α (purple) were computed per cell; left shows all coefficients, right shows those with p-value < 0.05. Percentages indicate the fraction of cells within each group that had a Pearson coefficient < 0; shown at the bottom. n >90 dHL60 cells (mEmerald-H3) and >110 cells (mEmerald-HP1α) from 4 replicates were analyzed.

Visualization and quantification of the time lapse movies of mEmerald-H3 transfected dHL60 cells undergoing NETosis revealed that H3 remains heterogeneously distributed inside the nucleus and a portion of H3 leaves the nucleus prior to nuclear rupture (**Figure 3A, C, D, F-H**). To assess the changes in the distribution of nuclear mEmerald-H3 during NETosis, we used the same metric as for DNA/chromatin homogenization (SD/MFI) and measured the SD/MFI of nuclear mEmerald-H3 at NETosis onset (@NO; assessed by MV shedding or cell rounding) and one minute before nuclear rupture (@NR; assessed by DNA release in the cytosol). We found that the ratio of SD/MFI remains close to 1 for mEmerald-H3 (SD/MFI^@NR^ : SD/MFI^@NO^ = 0.98 +/- 0.19 and < 1 in 51% of cells, **Figure 3C),** indicating that contrary to DNA, (SD/MFI^@NR^ : SD/MFI^@NO^ = 0.73 +/- 0.2 and < 1 in 92% of cells for DNA, **Figure S3C, H**), nuclear H3 does not homogenize during NETosis. This suggests that H3 maintains a specific, heterogenous pattern on accessible chromatin, which is consistent with the low mobile fraction of nuclear mEmerald- H3 in NETing cells (**Figure 2K**). To determine if the heterogenous distribution and slight but significant decrease in the mobile fraction of nuclear mEmerald-H3 was due to dissociated H3 exiting the nucleus before nuclear rupture, we quantified the mean fluorescence intensity (MFI) of nuclear and cytosolic mEmerald-H3 as well as the cytosolic to nuclear ratio of mEmerald-H3 as cells progress toward NETosis (**Figure 3D, F-H** and **S7A-C).** This revealed that mEmerald- H3 MFI decreases in the nucleus but increases in the cytosol between NETosis onset and nuclear rupture (**Figure 3 A, D, F-H**). We found a ∼ 16% decrease in the MFI of nuclear mEmerald-H3 (**Figure 3F**) and a ∼36% increase in the cytosolic to nuclear ratio of mEmerald- H3 (**Figure 3H**) between NETosis onset and nuclear rupture. Analysis of the peak of cytosolic mEmerald-H3 revealed a ∼17% increase in the maximum cytosolic mEmerald-H3 (**Figure 3G**). mEmerald-H3 leaks outside of the cytosol after plasma membrane permeabilization (drop in cytosolic intensity right before nuclear rupture in **Figure 3A, D**), explaining the non-detectable change in cytosolic mEmerald-H3 at nuclear rupture (**Figure 3G**) (30). Plotting the nuclear and cytosolic mEmerald-H3 at single cell resolution showed that the decrease in nuclear H3 coincides with the increase in cytosolic H3 (**Figure 3D**). Consistently, Pearson correlation analysis indicates that in 80.9 % of cells, the change in nuclear mEmerald-H3 negatively correlates with the change in cytosolic mEmerald-H3 (**Figure 3I**). Together, this data indicates that a portion of mEmerald-H3, ∼16%, translocates from the nucleus to the cytosol between NETosis onset and nuclear rupture, suggesting that dissociated nuclear H3 might become soluble cytosolic proteins.

To determine the nuclear and cytoplasmic dynamics of HP1α during NETosis, we analyzed the time lapse movies of mEmerald-HP1α transfected dHL60 cells undergoing NETosis and showed that HP1α homogenized before leaving the nucleus prior to nuclear rupture (**Figure 3B, C, E-H** and **S7D-F**). To assess the changes in the distribution of mEmerald- HP1α during NETosis, we compared the SD/MFI of mEmerald-HP1α between NETosis onset (@NO) and one minute before nuclear rupture (@NR). We found that SD/MFI^@NR^ : SD/MFI^@NO^ = 0.81 +/- 0.24 and < 1 in 81% of cells, (**Figure 3C**), indicating that HP1α homogenized between NETosis onset and nuclear rupture, although to a lesser extent than DNA (**Figure S3H**). To determine if the ∼40% decrease in the mobile fraction of HP1α during NETosis is due to HP1α exiting the nucleus, we measured the nuclear, cytosolic and cytosolic to nuclear ratio of mEmerald-HP1α between NETosis onset and one minute before nuclear rupture. We found that nuclear mEmerald-HP1α decreases in 71% of cells while peak cytosolic mEmerald-HP1α increases in 79% of cells. Population analysis indicates a ∼28% decrease in nuclear mEmerald- HP1α and ∼20% increase in the maximum cytosolic mEmerald-HP1α intensity (**Figure 3F, H**). Single cell analysis shows that the decrease in mEmerald-HP1α in the nucleus corresponds to its increase in the cytosol (**Figure 3B, E**). Consistently, Pearson correlation analysis showed a negative correlation between the changes in nuclear and cytosolic mEmerald-HP1α in 81.1% of cells (**Figure 3I**). 90% of the cells with significant Pearson coefficient (p-value < 0.05), show a negative correlation between the nuclear and cytosolic signal, further supporting that the decrease in nuclear mEmerald-HP1α coincides with the increase in the cytosolic pool. Together, this data indicates that a portion of mEmerald-HP1α (∼28%) translocate from the nucleus to the cytosol between NETosis onset and nuclear rupture, suggesting that dissociated nuclear HP1α might become soluble cytosolic proteins.

To determine if the distribution and location of mEmerald-HP1α mirrors the dynamics of endogenous HP1α, we used immunostaining to assess the distribution of endogenous HP1α during NETosis (**Figure S8,** see Materials and Methods). dHL60 cells were fixed 60 minutes after NETosis stimulation. Fixed cells were immunostained with an antibody against HP1α, along with LBR to monitor nuclear envelope integrity and identify stages of NETosis. High resolution 3D Z-stack of cells were acquired using a spinning disc confocal microscope.

Visualization (**Figure S8A**) and quantification (**Figure S8B-D**) of the fluorescent signal revealed that endogenous HP1α decreases in the nucleus prior to nuclear rupture. Analysis of the mean fluorescence intensity (MFI) of HP1α in relation to nuclear shape and integrity in NETing cells revealed a 2- and 4-fold decrease of nuclear HP1α MFI between NETosis onset and nuclear rounding and rupture, respectively (**Figure S8B**). Correlation analysis between HP1α MFI and the DNA SD/MFI showed a significant positive correlation further demonstrating that, similar to mEmerald-HP1α, endogenous HP1α translocates from the nucleus to the cytosol as chromatin homogenizes and increases in accessibility prior to nuclear rupture during NETosis (**Figure S8D**).

This data, together with the measured increase in the mobile fraction of H1, shows that chromatin structural proteins H1, H3 and HP1α dissociate from chromatin as chromatin accessibility increases during NETosis but while dissociated H1 remains inside the nucleus, dissociated H3 and HP1α leave the nucleus and become soluble cytosolic proteins.

We hypothesize that these dissociated chromatin binding proteins, will become macromolecular osmolytes which will increase the osmotic pressure in the cytosol and alter the mechanical properties of the plasma membrane (**Figure 4A**). We set out to test this hypothesis.

**Figure 4:**
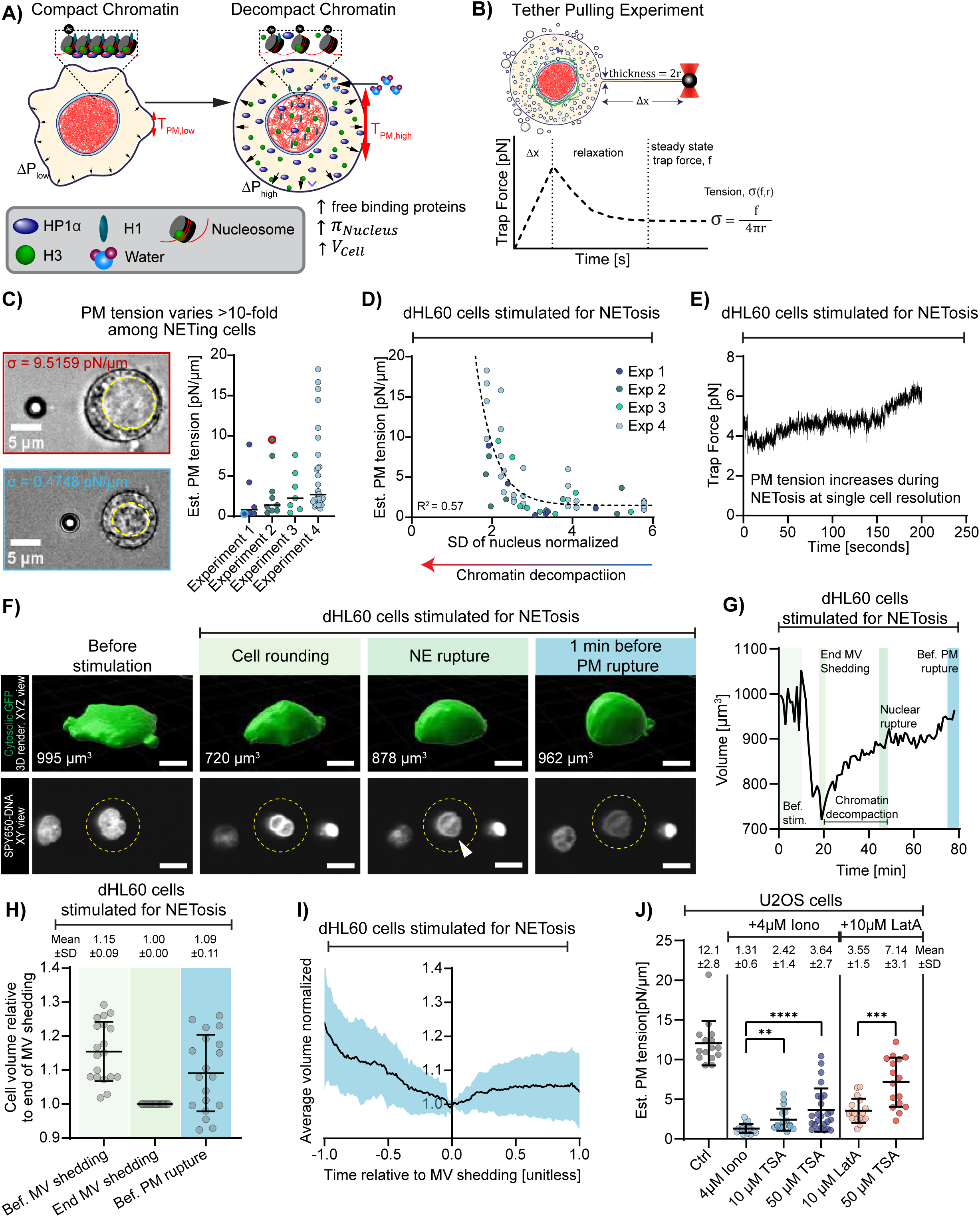
Chromatin decompaction increases cell volume and plasma membrane tension. **(A)** Working model of how chromatin decompaction (transition from compact to decompact chromatin) solubilizes and releases chromatin binding proteins in the cytosol which increases intracellular osmotic pressure (ΔP), cell volume and plasma membrane tension (T_PM_), disrupting cell integrity. **(B)** Schematic of a tether-pulling experiment (top) expected viscoelastic response (bottom), and estimation of membrane tension (σ in piconewton; pN/µm) from steady-state trap force (f) and tether thickness (r). **(C)** Representative images of NETosis-stimulated dHL60 cells attached to a bead (bright particle at the left of the image) via a membrane tether. Cells have completed microvesicle (MV) shedding and nuclear rounding but still have an intact looking nucleus outlined with a yellow dashed circle. Box around images to indicate cells on the right graph showing quantification of plasma membrane tension in a population of NETosis- stimulated dHL60 cells, across multiple (4) experimental replicates. Each data point represents a tether from a single cell. n = 29 NETosis-stimulated dHL60 cells, prior to nuclear rupture. **(D)** Correlation analysis between estimated (Est.) plasma membrane tension and chromatin decompaction (measured as normalized standard deviation (SD) of the nucleus in the brightfield SD channel; lower SD of nucleus = chromatin decompaction). Data were fitted with a one-phase decay nonlinear regression (dashed curve). Arrow on the bottom indicates a lower SD of the nucleus corresponds to more chromatin decompaction. **(E)** Representative single-cell trajectory of trap force over time. **(F)** Representative montage from a 3D lattice light sheet time-lapse movie of a dHL60 expressing cytosolic GFP (green; top) and stained with a DNA dye (SPY650- DNA; grey; bottom), stimulated for NETosis (ionomycin; 4 µM). Top: 3D rendering and XYZ view of the cell. Bottom, 2D, XY cross-section of the nucleus (SPY650-DNA). Raw cell volume is shown on top images. Scale bar, 10 μm. White arrowhead in bottom images to indicate nuclear rupture point (expansion of DNA into the cytosol). Yellow dashed circle indicates the nucleus corresponding to the dHL60 cell of interest shown above. **(G)** Quantification of raw cell volume for the NETing cell represented in **(F)**, from before NETosis stimulation to right before plasma membrane rupture. Shaded regions indicate different NETosis stages. Brackets in **(D-I)**, indicate the presence of ionomycin. **(H)** Quantification of normalized cell volume at three NETosis stages, relative to microvesicle (MV) shedding. Mean ± SD shown above graph. Each data point represents a cell **(I)** Mean ± SD of single-cell volume trajectories temporally aligned and normalized relative to microvesicle (MV) shedding (n = 19 cells, 3 replicates). **(J)** Quantification of estimated (Est.) plasma membrane (PM) tension in control (ctrl), ionomycin (Iono) +/- trischostatin A (TSA; a histone deacetylase inhibitor) or LatrunculinA (LatA; actin depolymerizing drug) +/- TSA treated U2OS cells. Mean ± SD shown at the top. n = 15, 21, 17, 25, 17, 16 cells per condition, respectively. Brackets denote the cell line and treatment. Statistical tests: Mann– Whitney U (two-tailed, nonparametric); ns: not significant; **** : p-value < 0.0001. All analyses were performed in GraphPad Prism v10.

### Chromatin decompaction inside the nucleus increases cell volume and plasma membrane tension during NETosis

To determine whether chromatin reorganization (homogenization and dissociation of chromatin binding proteins; hereafter referred to a “chromatin decompaction”) during NETosis and the resulting increase in soluble cytosolic proteins alters the mechanical properties of cells, we measured the tension at the plasma membrane as cells progress towards NETosis (**Figure 4, S9**). To measure plasma membrane tension, we used optical tweezers and performed classical membrane tether pulling experiments (61) on dHL60 cells undergoing NETosis (**Figure 4B, Movie S9**, see Materials and Methods). Cells were stained with CellMask (1 µM) to visualize the cell membrane and measure the diameter of the membrane tether. We used 2 µm carboxyl latex beads coated with concavalin A (1 mg/mL) to pull membrane tethers on NETing cells that have completed microvesicle (MV) shedding – indication of NETosis onset – but have not ruptured their nuclear membrane. This ensures that any measured change in plasma membrane tension is not due to chromatin directly interacting with or pushing against the cell membrane. From the trap force (**Figure S9A**) and the tether diameter (**Figure S9B**), we calculated the bending modulus and the estimated plasma membrane tension of each cell (**Figure 4C, D**, see Materials and Methods). We found that at the population level, the plasma membrane tension of NETing cells ranges from 0.5 pN/µm to 20 pN/µm (**Figure 4C)**. We observed that in cells with high plasma membrane tension, the signal of the nucleus in the brightfield channel appeared homogenous, suggestive of chromatin decompaction (**Figure 1E**, **Figure 4C-top**). Conversely, in cells with low plasma membrane tension, the chromatin appeared compacted – the nuclear signal in the brightfield channel was heterogenous (**Figure 1E**, **Figure 4C-bottom**). Plotting plasma membrane tension as a function of the standard deviation of the nuclear signal in the brightfield channel (BF SD) of NETing cells revealed a strong correlation between chromatin decompaction and plasma membrane tension (**Figure 4D**). Further, single cell measurement of plasma membrane tension over time (**Figure 4E**; **Movie S10**; see Materials and Methods) showed that membrane tension increases over time in NETing cells with a decompacted nucleus. This data indicates that chromatin decompaction during NETosis correlates with an increase in plasma membrane tension even while chromatin is still contained inside the nucleus.

### The increase in plasma membrane tension during NETosis is independent of the actin cytoskeleton and of membrane reservoirs

To determine the biophysical mechanism that mediates the observed increase in plasma membrane tension following chromatin decompaction during NETosis, we assessed the impact of known regulators of plasma membrane tension. Specifically, we tested the potential role of the actin cytoskeleton (62) and plasma membrane reservoirs (17) on plasma membrane tension during NETosis. We have previously shown (30) and confirmed here (**Figure S1**) that the actin cortex, which regulates plasma membrane tension, irreversibly disassembles at the early stages of NETosis, before the onset of microvesicle shedding. Since the increase in plasma membrane tension occurs after microvesicle shedding, this data indicates that the increase in plasma membrane tension during NETosis is independent of the actin cytoskeleton.

To assess the impact of plasma membrane reservoirs on plasma membrane tension during NETosis, we measured plasma membrane tension of single NETing cells as function of the length of the membrane tether (10, 20, 30, 40 µm; **Figure S9C**). We reasoned that if the plasma membrane reservoir decreased, the longer the membrane tether, the higher the membrane tension measured. Consistent with this reasoning, we found that plasma membrane tension of cells with decompacted chromatin increases with tether length (**Figure S9D**), suggesting that cells with decompacted chromatin have lower membrane reservoir. However, for each tether length, the plasma membrane tension of cells with decompacted chromatin (low SD of nucleus) is higher than that of cells with compacted chromatin (high SD of nucleus; **Figure S9C**). This result shows that the increase in plasma membrane tension with chromatin decompaction during NETosis (**Figure 4D**) is independent of plasma membrane reservoirs.

### Cell volume increases with chromatin decompaction during NETosis

To start probing whether chromatin decompaction inside the nucleus increases plasma membrane tension by increasing cellular osmotic pressure, we measured cell volume during NETosis in relation to chromatin decompaction (**Figure 4 F-I**). We reasoned that an increase in cellular osmotic pressure would translate into an increase in cell volume. dHL60 cells were transfected with cytosolic GFP (pmaxGFP) to stain the cytosol. Transfected cells were left to recover for 4 hours before being stained with a DNA dye (SPY650-DNA) to assess the different stages of NETosis (e.g., nuclear rounding, nuclear rupture). We used lattice light sheet microscopy to acquire 4 hours long, 3D time lapse movies of cells undergoing NETosis. Live cells were stimulated for NETosis 5 minutes after the onset of imaging to visualize the transition into NETosis. From the fluorescent intensity of cytosolic GFP, we generated masks of the cell body (minus microvesicles, **Figure 4F, Movie S11**) and calculated cell volume as cells progress toward NETosis (see Material and Methods). Quantification showed that cell volume rapidly (within 2 minutes) decreased as cells round up after microvesicle shedding (**Figure 4F-I**), potentially due to loss of cytosol after microvesicle shedding. But post microvesicle shedding, the cell volume persistently increases (up to 20 % in certain cells and relative to the volume at the end of microvesicle shedding) until the rupture or permeabilization of the plasma membrane and loss of GFP signal (**Figure 4G-I**). Further, the increase in cell volume during NETosis coincides with chromatin decompaction in NETing cells with intact nuclear membrane (**Figure 4F, G**). This data supports the hypothesis that chromatin decompaction and the resulting dissociation of chromatin binding proteins leads to an increase in cell volume and subsequent increase in plasma membrane tension.

### Chromatin decompaction inside the nucleus increases cell volume and plasma membrane tension in non-immune cells independently of NETosis

Our data so far shows that as chromatin decompacts inside the nucleus during NETosis (**Figure 1)** and chromatin binding proteins dissociate from chromatin (**Figure 2 and 3**), the cell volume and plasma membrane tension increase. This establishes a correlative relationship between chromatin decompaction, plasma membrane tension and overall mechanical state of the cell.

To evaluate the causal relationship between chromatin decompaction and plasma membrane tension, we set out to test it in non-immune cells outside of the context of NETosis. We choose the human osteosarcoma cell line U2OS cells as our model system because these cells do not possess the molecular pathways known to drive NETosis (**Figure S9E**; data from the “Human Protein Atlas” (63)). U2OS cells were treated with nothing (control) or different concentrations of Trichostatin A (TSA; 10 µM, 50 µM), a well-characterized histone deacetylase inhibitor that induces chromatin decompaction (64). To probe the impact of chromatin decompaction on plasma membrane tension, independently of the cytoskeleton, we treated U2OS cells with ionomycin (4 µM) – which depolymerizes the three cytoskeletal networks (65) - or with latrunculin A (10 µM) – which depolymerizes the actin cytoskeleton. When we used TSA, we pretreated cells with TSA for 1 hour and used optical tweezers to measure plasma membrane tension during the subsequent hour to minimize impact of gene expression.

Quantification showed that, consistent with the literature (66), disrupting the cytoskeleton drastically decreases plasma membrane tension (**Figure 4J**; membrane tension control: 12.1±2.8 pN/µm; ionomycin: 1.3±0.6 pN/µm; latrunculin treated: 3.55±1.5 pN/µm). Comparing plasma membrane tension as function of TSA treatment revealed that plasma membrane tension increased following TSA treatment of U2OS cells with disrupted cytoskeleton (**Figure 4J**). Notably, the increase in plasma membrane tension was dependent on the concentration of TSA (∼2- and 3-fold increase in 10 and 50 µM TSA treated cells, respectively, relative to ionomycin treated cells). Additionally, TSA treatment increased plasma membrane tension in cells treated with latrunculin A (2-fold increase) demonstrating that chromatin decompaction increased plasma membrane tension independently of the actin cytoskeleton. Together, this data establishes a causal relationship between chromatin decompaction and the increase in plasma membrane tension. It further shows that the impact of chromatin decompaction on plasma membrane tension is a general process, not limited to NETosis and its specific chromatin decompaction or other molecular pathways.

## Discussion

In summary, our work demonstrates that chromatin decompaction in the nucleus actively regulates the mechanical properties and state of the cell, independently of gene expression and of the cytoskeleton. We found that during NETosis, systemic chromatin decompaction starts before nuclear rupture and correlates with increased plasma membrane tension. We showed that chromatin decompaction during NETosis occurs via dissociation of major chromatin binding proteins that regulate chromatin structure (**Figure 2, 3**), resulting in a progressive increase in chromatin accessibility as cells progress toward different stages of NETosis (**Figure 1**). For instance, we found that the linker histone H1, the nucleosomal histone H3 and the heterochromatin binding protein HP1α dissociate from chromatin during NETosis (**Figure 2, 3**). We showed that dissociated chromatin structural proteins H3 and HP1α translocate from the nucleus to the cytoplasm (**Figure 3**), where we posit, they become osmolytes and increase osmotic pressure. Consistently, we found that chromatin decompaction during NETosis correlates with an increase in cell volume and plasma membrane tension (**Figure 4 C-I**). We demonstrated the causal relationship between chromatin decompaction and plasma membrane tension by showing that inducing chromatin decompaction in non-immune, U2OS cells with disrupted cytoskeleton resulted in an increase in plasma membrane tension (**Figure 4J**).

Together, this data supports our initial hypothesis that chromatin can actively regulate the mechanical state of cells independently of the cytoskeleton, establishing chromatin as a long-range/contact-less force generator.

Our work demonstrates that plasma membrane tension and volume increases prior to plasma membrane rupture during NETosis providing a new framework for understanding how the plasma membrane ruptures during NETosis. The culmination of suicidal NETosis is the rupture of the plasma membrane to allow chromatin to reach the extracellular environment (30, 41, 67). Plasma membrane rupture can be driven by biochemical modifications that weaken the membrane (68), by physical forces that overcome the lysis tension of the membrane (69) or a combination of both. Our measured increase in plasma membrane tension during NETosis supports the model where rupture of the plasma membrane during NETosis is driven by physical forces applied on the membrane resulting in an increase in membrane tension. Cells will rupture when the applied forces are greater than the lysis tension of the plasma membrane. Our data supports previous work by Neubert et al. that showed that plasma membrane tension increases during the late stages of NETosis (41). The authors proposed that chromatin swelling in the cytosol after nuclear rupture, pushes against the plasma membrane applying entropic elastic forces to the plasma membrane. The novelty of our work lies in showing that plasma membrane tension increases during NETosis while chromatin is still located inside the nucleus (**Figure 4**), precluding the role of chromatin-generated entropic elastic forces. Rather, our data converge toward a model where chromatin decompaction in the nucleus by releasing chromatin binding proteins generates new osmolytes that increase the cellular osmotic pressure, volume then plasma membrane tension. Thus, a potential unifying model is that chromatin decompaction generates osmotic pressure or entropic elasticity on the plasma membrane depending on the location of chromatin. Further quantitative measurements of plasma membrane tension throughout NETosis, relative to chromatin decompaction in the nucleus and swelling in the cytosol, will be important to test this unifying model.

Our demonstration of the progressive increase in chromatin accessibility and dissociation of large portions of the linker histone H1 and the heterochromatin protein HP1α provides a new framework for understanding the molecular mechanism of chromatin decompaction during NETosis. Previous studies have focused on identifying the upstream regulators of chromatin decompaction during NETosis (35). For instance, PAD4 (70), neutrophils elastase (46), MPO (46) and Calpain (71) were shown to regulate chromatin decompaction during NETosis. However, how these upstream regulators results in chromatin decompaction during NETosis has remained unclear (65). Our data aligns with previous literature (35, 70) and demonstrates that the nucleosomal histone H3 is citrullinated at the early stages of NETosis (**Figure 2A, B, S4**). However, we found that only a small portion of H3 (∼16%) dissociates from chromatin during NETosis (**Figure 3A, F)**. Importantly, our data revealed that large portions of proteins that mediate higher order chromatin structures than nucleosome, H1 and HP1α, dissociate during NETosis. We showed that while H1 mobility increases to a “set” point in NETing cells, independently of the level of chromatin homogenization, HP1α progressively dissociates from chromatin as chromatin homogenizes during NETosis (**Figure S6B, C**). This suggests that H1 dissociation precedes that of HP1α supporting a model where chromatin decompaction during NETosis might be mediated by disconnecting nucleosomes through H1 dissociation, followed by progressive undoing of higher order chromatin compaction states such as chromatin loops and topology associated domains (72). This model is consistent with early finding that chromatin in NETs has the “beads on a string” configuration of nucleosome bound DNA (42). Important questions arise from our work. First, what is the molecular mechanism of H1 and HP1α dissociation from chromatin during NETosis? Is it through citrullination or via other post translational modifications such as phosphorylation known to regulate H1- or HP1α- chromatin interaction (73, 74)? Second, what are the dynamics of other chromatin structural proteins or epigenetic readers that recognize open chromatin? Third, does H3 citrullination participate in chromatin decompaction, not by dissociating H3 from chromatin but by destabilizing the interaction between nucleosomes and their structural proteins? Note that our data does not exclude the possibility that a large fraction of nucleosomes dissociate from chromatin during NETosis, but we could not detect such dissociation events due to the larger size and corresponding lower mobility of nucleosomes in the confined nucleus. Further molecular, biophysics and genetics works will be important to identify the sequence of events leading to chromatin decompaction during NETosis and determine whether and how posttranslational modifications of histones and other chromatin binding proteins participate in chromatin decompaction during NETosis.

Beyond NETosis, the importance of our finding lies in demonstrating that mammalian cells have an alternative mechanism of increasing plasma membrane tension during physiological processes – NETosis – and this mechanism does not use the cytoskeleton. The prevailing model in the field is that the cytoskeleton which applies force on the membrane and regulates membrane composition and membrane reservoir, “sets the value” of plasma membrane tension (75, 76). Consistent with this model, we showed that plasma membrane tension drastically decreases in cells with disrupted cytoskeleton (**Figure 4J**). However, our measurements in NETosis and cancer cells with decompacted chromatin demonstrated that the tension of a mammalian cell membrane devoid of or with weakened cytoskeleton can be as high as 20 pN/µm, mirroring the tension of mammalian cells with intact cytoskeleton (**Figure 4I**, (77)).

What are the molecular and biophysical mechanisms of this tension increase? Besides the cytoskeleton, plasma membrane reservoirs, membrane trafficking and osmotic pressure can regulate plasma membrane tension (76, 78). We showed that the increase in plasma membrane tension during NETosis is independent of membrane reservoir and correlates with increased cell volume. Membrane trafficking requires the cytoskeleton and ATP which are disassembled and depleted, respectively, at the early stages of NETosis (30, 41) prior to membrane tension increase. The correlation between membrane tension, cell volume and increased mobility of chromatin binding proteins, supports the hypothesis that chromatin decompaction increases osmotic pressure. However, our data does not exclude the possibility that other lipid enzymes or trafficking machinery might be activated by chromatin decompaction or the loss of the cytoskeleton to alter the composition and mechanical properties of the plasma membrane.

Future lipidomic and surface proteomics studies combined with membrane biophysics studies will be important to determine whether and how disrupting the cytoskeleton and decompacting chromatin increases plasma membrane tension by altering the molecular composition of the plasma membrane.

Finally, our demonstration that chromatin decompaction changes the mechanical state of the cell provides a new framework for understanding the role of chromatin in mammalian cell biology. While our study focused on neutrophil physiology – NETosis – and generalized our findings in a model cancer cell line, data from the literature indicate that chromatin compaction state might regulate the mechanical properties of cells in other physiological process (79, 80).

For instance, Zlotek-Zlotkiewicz et al. and Son et al., demonstrated that during mitotic entry - the transition between G2 and mitosis when chromatin drastically reorganizes into chromosomes - the volume of the cell increases by 10 to 20%, independently of the actin cytoskeleton and endocytosis but due to water influx across the plasma membrane (79–81). Mathematical modelling by Romain et al, proposed that this osmotically driven increase in cell volume at mitotic entry is due to changes in chromatin compaction state, in particular increase in histone acetylation (82). Consistently, our data show that decompacting chromatin through histone acetylation – with TSA – increases plasma membrane tension. Importantly, our data with NETosis, where histone acetylation has minimal roles (83), indicates that the impact of chromatin decompaction on cell mechanics is not specific to histone acetylation. Importantly, both chromatin compaction during mitosis (79, 80, 82) and decompaction during NETosis results in an increase in cell volume suggesting that any drastic variation of chromatin compaction state might alter the mechanical properties of cells. In the future, it will be important to determine whether and how the biophysical impact of chromatin reorganization on plasma membrane tension and cell volume regulates other biological processes such as cell migration, differentiation or immune cell activation during which mammalian cells reorganize chromatin and alter their mechanical properties.

Together, our data demonstrates that chromatin decompaction inside the nucleus can directly regulate cell volume and plasma membrane tension independently of gene expression and the cytoskeleton. This establishes chromatin as a contactless force generator broadening our understanding of the biophysical role of chromatin in mammalian cells. Our work opens new avenues for understanding whether and how cells actively alter the compaction state of chromatin to accomplish essential functions including migration and differentiation.

## Acknowledgments

We thank members of the Thiam lab for discussions and providing feedback on the manuscript, Lacra Bintu for discussions, David P Lenzi from CSIF: Stanford University Cell Sciences Imaging Core Facility (RRID:SCR_017787). We thank the funding agencies who support our work. A.C is supported by the Stanford Graduate Fellowship. M.S and M.K were supported by the SoM Dean’s Postdoctoral Fellowship, Stanford, University. H.R.T and the Thiam lab is supported by the CZ Biohub San Francisco, the David and Lucile Packard Foundation, Stanford Bio-X, the Koret Foundation and the Esther Ehrman Lazard Faculty Scholar Award.

## Data availability statement

Data generated and code developed in this paper will be made available on data repositories and GitHub after the peer review process. Data for graphical representation of RNA transcript abundance in Figure S9E was obtained from the Human Protein Atlas, www.proteinatlas.org, (Uhlén M et al, 2015) and is available at the URL: v22.proteinatlas.org<u>/humancell</u>

## SUPPLEMENTARY MATERIALS

### MATERIALS AND METHODS

#### Cell culture

A human promyelocytic leukemia cell line, HL-60 cells, was purchased from ATCC (ATCC® CCL-240™), cultured in RPMI 1640 medium (Cytiva, SH30096.01) supplemented with 1% glutamax (Thermo Scientific, 35050061), 1% Penicillin/Streptomycin (Pen/Strep; Gibco, 15070063), 25 mM HEPES (Cytiva, SH30237.01) and 15% heat-inactivated Fetal Bovine Serum (FBS; R&D Systems, S11150) and split every 3 days at 2×10^5^ cells/mL. Cells were maintained at 37°C in a humidified 5% CO_2_ incubator. Cells were differentiated into dHL60 cells via treatment with culture media supplemented with 1.3% dimethyl sulfoxide (DMSO; Sigma-Aldrich D2650). Differentiated cells were used at day 6 or 7 after treatment with DMSO.

A human osteosarcoma cell line with epithelial morphology, U2OS cells, was purchased from ATCC (ATCC® HTB-96), and cultured in McCoy’s 5A Medium (Cytiva, SH30200.01) supplemented with 1% Penicillin/Streptomycin (Pen/Strep; Gibco, 15070063), and 10% Fetal Bovine Serum (FBS; R&D Systems, S11150). Cells were split every 3 days or at 80-90% confluency with trypsin-EDTA 0.25% (Thermo Scientific, 25200072) at 2×10^5^ cells in tissue culture treated 25 cm^2^ flask (CELLTREAT. 229331) and maintained at 37°C in a humidified 5% CO_2_ incubator.

#### Inhibitors

The following inhibitors were purchased from companies indicated in brackets: Cycloheximide (Selleckchem, S7418), Actinomycin D (Sigma-Aldrich, A9415), Trichostatin A (TSA; Tocris, 1406), Latrunculin A (Millapore Sigma, 428021).

#### cDNA expression vectors

cDNA encoding mKOet-H1 (Addgene, #57922), mEmerald-HP1α (Addgene, #54124), mEmerald-H3.3 (Addgene, #54116) were purchased from Addgene. pmaxGFP was purchased from Lonza (Lonza, VCA-1003).

#### Transient transfection of cDNA in dHL60 cells

1 mL of culture medium was prewarmed in a sterile 1.5 mL eppendorf tube and in a 24-well plate. A mixture of 100 μL Nucleofector® V Solution (Lonza, VCA-1003) containing endotoxin- free DNA (2μg for mKOet-H1; 3μg for mEmerald-HP1α; 1μg for mEmerald-H3.3 was equilibrated at room temperature for 5 min. 2×10^6^ dHL60 cells were pelleted at 2000 rpm for 5 min at room temperature, resuspended in the DNA+V Solution mix, and electroporated (Y-001 program, Amaxa® Nucleofector®). Immediately, 1 mL of prewarmed medium from eppendorf tube was added to the cuvette, and the suspension was transferred back to the eppendorf tube and incubated for 30 min at 37 °C before transferring to the pre-equilibrated well of a 24-well plate. Cells were imaged at least 10 hours after transfection.

#### Transient transfection of cDNA in U2OS cells

1 mL of complete medium was prewarmed in glass bottom 35 mm culture dish (FluoroDish, FD35-100). A mixture of 100 μL cytoskeleton buffer (CB; 10 mM MES, 138 mM KCl, 3 mM MgCl2, 2mM EGTA) and 1 μg of pmaxGFP^TM^ (Lonza, VCA-1003) was equilibrated at room temperature for 5 min. 5×10^5^ U2OS cells were pelleted at 130xG for 5 min at room temperature, resuspended in the DNA-cytoskeleton buffer solution mix, and electroporated (X-001 program, Amaxa® Nucleofector®). Immediately, 75k cells were transferred into the 35 mm culture dish and left to recover overnight.

#### Preparing cells for live imaging

*<u>Preparing dHL60 cells for NETosis assay via live cell imaging</u>*: dHL60s non transfected or expressing fluorescently tagged proteins, were stained with 1 µM of SPY650-DNA dye (Cytoskeleton, CY-SC501), and/or CellMask (Thermo Scientific, C10045), and/or SPY555- FastAct (Cytoskeleton, CY-SC205) for 1 hr at 37°C in a humidified 5% CO_2_ incubator. Cells were then pelleted down at 2000 rpm for 5 minutes at room temperature and resuspended in imaging media [serum-free, no phenol red RPMI 1640 medium (Gibco, 11835030) buffered with 25 mM HEPES, 1% Pen/Strep, 1x Glutamax]. 10^5^ to 3×10^5^ were used for live imaging.

*<u>Preparing dHL60 and U2OS cells for optical tweezer experiments:</u>* Glass coverslides and coverslips (#1.5) were washed with ethanol (70%) and water prior to experiment. Day 6 or day 7 dHL60 cells were stained with CellMask (1:1000) at 8×10^5^ cells/mL for 20 minutes. 250 µL of the cell suspension was spun at 2000 rpm and resuspended in 250 µL of imaging media. Carboxyl latex beads (2 μm, Invitrogen, C37278) were coated for 1 hour in 1mg/mL concanavalin A (Millipore sigma, C2010) and added to cells at 10 beads per cell. Double sided adhesive tape (Duo Double Sided Adhesive Tape, 75600600) was folded to create a single layer approximately 200 μm thick, then attached to the coverslip. Cells and beads were first deposited on the coverslip for 5 minutes to allow dHL60 cells to adhere before addition of ionomycin (4 µM final concentration). The total volume was 15-20 µL. A coverslide was used to seal the chamber which was mounted on the optical tweezer for imaging.

#### Cell fixation and permeabilization

5×10^4^ dHL60 cells in 20 µL of imaging media were seeded on plasma cleaned 12 mm diameter #1.5 glass coverslips (EMS Cat. #72290-04). Seeded cells were given 5-10 minutes to adhere on coverslips at 37°C in a humidified 5% CO_2_ incubator. For stimulated cells, an additional 200 µL of imaging media with ionomycin calcium salt (Sigma-Aldrich, I0634; final concentration of 4 μM) was added to the coverslip and cells were incubated for 1 hour at 37°C in a humidified 5% CO_2_ incubator. Unstimulated cells were fixed immediately after the initial 5-10 minutes. Cells were fixed with 4% paraformaldehyde (PFA; Electron Microscopy Science, 15710) in 1x cytoskeleton buffer (CB; 10 mM MES, 138 mM KCl, 3 mM MgCl2, 2mM EGTA) for 20 minutes at 37°C, unreacted PFA was quenched with glycine (0.1 M glycine in 1x CB), then permeabilized with 0.5% Triton-X100 in 1x CB for 10 minutes at 37°C. Cells were washed with washing buffer containing 0.1% Tween20 (Sigma-Aldrich, P7949) in 1x Tris-buffered saline (TBS, 20 mM Tris- HCl (pH 7.6), 150 mM NaCl_2_) two times at room temperature for 5 minutes.

*<u>Indirect immunofluorescence:</u>* Formaldehyde-fixed and triton-permeabilized cells were incubated in blocking solution (2% BSA; 0.1% Tween20 in 1x TBS) for 1 hour at room temperature. Cells were stained for 2 hours at RT with primary antibodies diluted in blocking solution. Cells were then washed (3 x 5 minutes) in washing buffer (0.1% Tween20 in 1x TBS). After the wash, cells were incubated with fluorophore-conjugated secondary antibodies and DAPI (1ug/mL; Sigma-Aldrich D9542) diluted in blocking solution, for 1 hour at room temperature. Finally, cells were washed (3 x 5 minutes) in washing buffer before being mounted on a glass slide in mounting media (Dako fluorescence mounting media, S302380-2). Primary antibodies used: mouse anti-LBR antibody (Abcam, BBmLBR 12.F8; dilution 1:500), rabbit anti-histone H3 (citrulline R2+R8+R17) antibody (Abcam, ab5103; 1:1000), rabbit anti-CBX5 (HP1α) antibody (Sigma-Aldrich, HPA016699; dilution 1:200). The secondary antibodies used: Alexa Fluor 488 Goat anti-rabbit (Thermo Fisher Scientific, A-11008; dilution 1:400), Alexa Fluor 594 Donkey anti-mouse (Jackson ImmunoResearch, 715-587-003; dilution 1:500)

*ATAC-SEE*:

*<u>Tn5 transposase assembly:</u>* Recombinant purified hyperactive Tn5 transposase loaded with Alexa488 fluorophore-conjugated adaptors (Alexa488-Tn5) were kindly provided by our collaborator Xie Liangqi (Lerner Research Institute, Cleveland Clinic, Ohio, USA). The adaptor sequences are as follows:

Tn5ME-A-Alexa488: 5′-Alexa488-TCGTCGGCAGCGTCAGATGTGTATAAGAGACAG-3′; Tn5ME-B-Alexa488: 5′-Alexa488-GTCTCGTGGGCTCGGAGATGTGTATAAGAGACAG-3′; Tn5MErev: 5′-(phos) CTGTCTCTTATACACATCT-3′.

*<u>Tn5 tagmentation:</u>* An aliquot of 2x Tagment DNA buffer (TD; 10 mL of 2x TD buffer, mix 200 µL of 1 M Tris-HCl (pH7.6), 200 µL of 0.5 M MgCl_2_, 2 mL of DMF (N,N-Dimethylformamide; Sigma-Aldrich D4551), was diluted in 7.6 mL of sterilized dH20. Formaldehyde-fixed and triton-permeabilized cells were incubated in tagmentation solution (100 nM Alexa488-Tn5 in 1X TD buffer) overnight at 4°C. Cells were then washed with freshly made ATAC-see washing buffer (0.01% SDS, 50 mM EDTA in 1x dPBS) three times for 15 minutes at 55°C. To label the nuclear envelope and DNA, immunostaining was performed for Lamin B Receptor (LBR) and cells were stained with DAPI and SPY650-DNA as described in the indirect immunofluorescence section of the Methods. Coverslips were mounted on a glass slide in mounting media (Dako fluorescence mounting media, S302380-2). The samples were imaged on a spinning disc confocal microscope. 3D Z stack images were acquired at xx interval covering the entire cell.

#### Microscopy

*<u>Spinning disc confocal and DIC microscopy:</u>* Imaging was performed on a Nikon Eclipse Ti2 inverted microscope equipped with Perfect Focus, an Okolab stage-top incubator for controlled temperature, humidity and CO_2_, a Crest X-light V3 spinning disc scanhead, a Kinetix sCMOS camera, a Plan Apo 60× oil 1.4 NA DIC Nikon objective lens, a Nikon LUNF (405nm, 90mW) and Nikon Opti-Microscan-01 for FRAP/Photostimulation, and the appropriate DIC prisms in place. Illumination was provided on the Crest V3 by a 7-line solid-state Celesta Laser Unit, 1 W per channel (405 nm; 445 nm; 488 nm; 515 nm; 561 nm; 640 nm; and 730 nm power measurement is at the fiber output). DIC illumination was provided by an LED. Microscope was equipped with the Nikon motorized stage with xy linear encoders and a closed-loop *Z*-axis Piezo nanopositioning system with 200 μm travel. Laser confocal or DIC illumination was selected with electronic shutters and an automated filter turret containing a multibandpass dichromatic mirror together with an electronic emission filterwheel. Microscope functions were controlled with NIS- Elements software (Nikon).

For NETosis assay by live imaging, a non-coated, gamma-irradiated glass bottom 35 mm dish (World Precision Instruments, FD35-100) or 4 Well chambered cover Glass (CellVis, C4-1.5H- N) was placed on a pre-warmed (37°C) microscope stage. Sample temperature, humidity and CO_2_ level were controlled with the respective microscope’s stage-top incubator. Cells (10^5^ to 3×10^5^ cells) were plated on temperature equilibrated glass-bottom dishes for 5 minutes.

Random fields (3–10) containing multiple adherent cells were selected for imaging. Confocal, DIC or brightfield images at the coverslip-cell interface and 3 μm above in the cell center were acquired for each position. The image acquisition was paused 3-5 minutes after beginning, ionomycin calcium salt (Sigma-Aldrich, I0634; final concentration of 4 μM) was added and imaging was resumed. Images were acquired every 1 minute for 4 hours, at maximum 5% laser power.

Fixed dHL60 cells immunostained with anti-LBR, anti-histone H3 (citrulline Arg2+Arg8+Arg17) and anti-CBX5 (HP1α) antibodies as well as ATAC-see samples mounted to glass slides were imaged by spinning disc confocal microscopy. For each coverslip, multiples (3–6) random fields were selected, and pairs of DIC and fluorescence confocal stacks were captured. For samples co-immunostained with Tn5-Alexa488 + anti-LBR + DAPI/SPY650-DNA, images were acquired with 488 nm, 561 nm and 405/640 nm lasers at 20%, 20% and 10%/50% laser power respectively. For samples co-immunostained with anti-histone H3 + anti-LBR + DAPI, images were acquired with 488 nm, 561 nm and 405 nm lasers at 1%, 50% and 15% respectively. For samples, co-immunostained with anti-CBX5 + anti-LBR + DAPI, images were acquired with 488 nm, 561 nm and 405 nm lasers at 10%, 50% and 20% laser power respectively.

*<u>Laser scanning microscopy:</u>* To generate the nuclei heatmap of live dHL60 cells, images were acquired on a Zeiss LSM 900 Airyscan laser scanning microscope equipped with a Plan- Apochromat 63×, 1.40 NA oil immersion objective. The pinhole was set to 1 Airy unit (56 µm), and illumination was provided using a 640 nm laser at 2% power. Microscope functions were controlled with the ZEN software (Zeiss).

*<u>Fluorescence recovery after photobleaching (FRAP) experiments:</u>* FRAP experiments of mEmerald-H3 and mKOet-H1 were performed on the spinning disc confocal microscope equipped with a FRAP module described above. Fluorescent proteins were bleached within a ∼3 μm^2^ region using a 405 nm laser at 10% laser power with a dwell time of 300 μs. Images were acquired every 5 seconds before and after bleaching using a triggered acquisition feature on NIS-Elements software where we imaged with 488 nm, 640 nm, and 561 nm lasers at 5% power and 200 ms exposure.

FRAP experiments of mEmerald-HP1α and mKOet-H1, were performed on a Nikon Ti2 inverted microscope equipped with Perfect Focus, Okolabs blackout enclosure with environmental control, a Crest X-Light V3 spinning disk, a KINETIC 10MP CMOS 25 mm camera, a Plan Apo 60× Oil 1.4 NA DIC Nikon objective lens, iLAS2 FRAP/Photostimulation (405/473 nm; 50/30 mW), and the appropriate DIC prism in place where illumination was provided on the Crest V3 by a 7-line solid state Celsta Laser Unit (∼800 mW/line; 365 nm, 440 nm, 488 nm, 514 nm, 561 nm, 640 nm, 730 nm). DIC illumination was provided by an LED and microscope was equipped with a motorized XY stage. Fluorescent proteins were bleached within a ∼3 μm^2^ region using a 473 nm laser at 20 % with a dwell time of 300 μs. Images were acquired every 5 seconds before and after bleaching using a triggered acquisition feature on NIS-Elements software where we imaged with 475 nm, 640 nm, and 549 nm lasers at 5% power and 200 ms exposure.

*<u>ZEISS Lattice Light Sheet 7 for volumetric measurements:</u>*Imaging was performed on a ZEISS Lattice Light Sheet 7 microscope equipped with an enclosed stage-top incubator (temperature, humidity and CO_2_ control), a Hamamtsu Orca sCMOS camera, a 13.3× water 0.4 NA illumination objective and a 44× water 1.0 NA imaging objective tilted 30 degrees with respect to the sample. The instrument was equipped with a motorized 5-axis stage, 488, 561, and 647 nm laser lines, and an LED for brightfield illumination. Before imaging, we calibrated the X and Y tilt, focused the sheet, focused the waist, and adjusted aberration control. We used a 100 μm x 1800 nm lightsheet and imaged dHL60 cells undergoing NETosis by illuminating cytosolic GFP and SPY650-DNA with the 488 nm and 647 nm lasers, respectively, at 0.7% for 33 ms using the LBF 405/488/561/642 emission filter. Image acquisition was paused 5 minutes after beginning, 4 μM ionomycin calcium salt was added, and imaging was resumed every 1 minute for 4 hours.

The tilted image sets were de-skewed using ZEN software to generate 3D stacks with conventional X/Y/Z coordinates.

*<u>Optical tweezers</u>:* The optical tweezer (OT) platform (SENSOCELL, IMPETUX OPTICS, Spain) consists of a continuous wave laser (λ = 1064 nm, 5-W nominal output power) steered with a pair of acousto-optic deflectors (AOD1 and AOD2) and a force detection module. This is mounted around Nikon Eclipse Ti2 inverted microscope equipped with Perfect Focus, a custom Nikon stage-top, and a Nikon D-LEDI. The laser is directed onto a microscope objective (Plan Apo 60× water immersion 1.2 NA Nikon objective lens) after entering through the epifluorescence port. To prevent IR light leaking toward the detector, a neutral, shortpass filter (IR-F) was placed at the imaging optical path. The optical traps were positioned using LightAce software (IMPETUX OPTICS, Spain), in synchronization with an ORCA-Flash4.0LT3 HAMAMATSU camera recording BF and fluorescent images of the sample. The force detection module is a collecting lens requiring immersion oil (1.4 NA) that operates by detecting light-momentum changes (85). The module enabled us to measure forces with no trap pre-calibration.

#### Image Analysis

*<u>Quantification of the percentage and timing of cellular events during NETosis:</u>* Time-lapse spinning disk confocal and differential interference contrast (DIC) movies of cells stained with SPY650-DNA were used to quantify the percentage and timing of NETosis events as previously described ref (30)*.<u>Quantification of the percentage and timing of cellular events during NETosis:</u>* Time-lapse spinning disk confocal and differential interference contrast (DIC) movies of cells stained with SPY650-DNA were used to quantify the percentage and timing of NETosis events as previously described ref (30). Events quantified : Plasma membrane microvesicle shedding, defined as the release of vesicles from the cell periphery as seen in DIC movies; chromatin reorganization, defined as the decrease in SPY650-DNA inhomogeneity as seen in confocal fluorescence movies; Nuclear rounding, defined as the establishment of a circular nuclear periphery as seen in DIC and confocal fluorescence color overlay movies; Nuclear rupture, defined as expansion of the DNA periphery outside of the nuclear boundaries into the cytosol as seen in DIC and confocal fluorescence color overlay movies; PM permeabilization, defined as a decrease in contrast at the cell periphery as seen in DIC movies; and NETosis, defined as expansion of DNA outside of the cell boundary as seen in DIC and confocal fluorescence color overlay movies. All adherent cells in imaging fields were included in the analysis of the percentage of cells undergoing an event while only cells that remained in the field for the entire duration of the movie were included in the quantification of the timing of events. Multiple imaging fields were analyzed for each condition until the number of cells per condition was equal or greater than 50. The timing of an event was defined as the first time point when the corresponding event was observed in the movies by eye.

*<u>Quantification of the standard deviation and roundness of the nucleus during NETosis:</u>* Time-lapse spinning disk confocal and brightfield (BF) movies of dHL60 cells stained with a DNA dye (SPY650-DNA) were used to quantify the change in DNA and nuclear homogeneity during NETosis. For each time point, individual NETing cells were cropped and analyzed with a custom python script that applied an intensity thresholding to segment individual nuclei (based on the DNA channel), from which the standard deviation and roundness were calculated in the SPY650-DNA and the brightfield channel with respect to time. Cells with poor nuclear mask were excluded from the quantification. Each NETing cell was labeled for the timing when NETosis onset and nuclear rupture or end of the movie (whichever happened first). Single cell trajectories (SD of DNA, SD of BF, roundness) were temporally (x-axis) aligned so that they all start at NETosis onset and end 1 minute before nuclear rupture / end of the movie. Amplitudes of single cell trajectories (SD of DNA, SD of BF, roundness) were subjected to a median filter and normalized relative to their values at NETosis onset enabling direct averaging and comparison across asynchronous cells.

*<u>Quantification of ATAC-see and protein of interest (POI) in fixed cells</u>:* Fixed-cell images (immunostained for Tn5, Pan-H3Cit, HP1α) were acquired as 16-bit multi-channel, multi-Z stacks on a spinning disc confocal microscope. For each cell, an analysis focal plane (Z_selected) was determined from the DNA channel (DAPI or SPY650-DNA). Nuclear masks were generated using histogram-mode thresholding with an added factor of *k·SD* (where *k* = 1.0–2.0, empirically optimized on pilot datasets and held constant thereafter). Small objects (<100 pixels) were excluded. Nuclear area was summed across slices, and the slice with the maximum summed area was designated as Z_selected. For quality control, Z_selected and its adjacent slices were visually inspected and adjusted if needed. Z_selected was used to generate 1) nuclear masks from the DNA channel using the same mode+*k·SD* thresholding, followed by removal of small objects and binary hole filling; 2) Images of POI (Tn5, Pan-H3Cit, HP1α) ensuring that all analysis is done on the same focal plane. To estimate background intensity, each POI image was Gaussian-blurred (5 × 5 kernel) to stabilize its histogram, and thresholded at (histogram mode + 1 gray level) to obtain a conservative foreground selection. Morphological opening, closing, and dilation further refined the foreground mask. Pixels outside this mask were defined as background. The mean intensity of background pixels was calculated from the original (non-blurred) 16-bit POI image and subtracted uniformly from all pixels.

Negative values were set to zero, and all values were clipped to the 16-bit range. All downstream analyses were performed on these background-corrected images. For each nucleus, integrated intensity of the POI was calculated as the mean background-corrected intensity within the nuclear mask multiplied by the nuclear area, equivalent to summing all pixel values within the mask. Nuclear areas were converted to µm² based on the microscope- reported pixel size. Image processing and quantitative analyses were performed using custom Python code (tifffile, NumPy, OpenCV, SciPy, scikit-image, pandas, and matplotlib).

*<u>Generation of the heatmap of nuclei in NETing cells</u>*: Single-plane images of NETosis-stimulated (ionomycin calcium salt, 4 µM) dHL60 cells stained with a DNA dye (SPY650-DNA) were used to generate average heatmaps of the nucleus at different stages of NETosis. Fields were tracked until the DNA of the same nucleus could be imaged in two stages: Early NETosis (bright peripheral rim with uneven intra-nuclear intensity) and Chromatin reorganization (loss of the rim and homogenized signal). Twenty nucleus pairs were collected (n = 20 cells). For each nucleus, a binary nuclear mask was generated from the DNA channel and corrected manually when necessary. The mean DNA intensity from the Early NETosis frame was used to normalize both images from that nucleus, such that values represent fold change relative to the early-phase mean. To average spatial patterns, nuclei were size-standardized: the masked nuclear crop was resized, centered on a 128 × 128 canvas, and multiplied by a circular mask whose radius matched the phase-specific mean radius measured across the dataset (Early NETosis: 37.5 px; Chromatin reorganization: 33.8 px; pixel size 0.098 µm·px⁻¹). Standardized, fold-normalized images were then averaged pixel-wise across cells to produce the heatmaps. Image processing and analysis were performed in Python using custom code.

*<u>Quantification of F-actin dynamics during NETosis:</u>* Time-lapse spinning disk confocal and DIC movies of dHL60 cells stained with a DNA dye (SPY650-DNA) and F-actin dye (SPY555- FactAct). Images at the cell-coverslip interface were acquired every 5 seconds for 30 minutes to monitor fast F-actin dynamics. Integrated intensity of the F-actin fluorescence signal, background corrected, was measured for each cell by manually tracing whole cells in 3 stages: Before NETosis onset (before ionomycin was added), Onset of NETosis (when cells were visually initiating NETosis (MV shedding or onset of rounding) and there was a peak in actin fluorescence), and actin disassembly (the last time point of the 30 minute movie). Fold change between each two states was computed and single cell trajectories were temporally aligned relative to MV shedding.

*<u>Quantification of FRAP measurements:</u>* Time-lapse spinning disk confocal movies were acquired before and after photobleaching dHL60 cells stained with a DNA dye (SPY650-DNA) and expressing mKOet-H1 and mEmerald-H3 or mEmerald-HP1α. For each time point, individual nuclei were cropped and analyzed with a custom python script that applied an intensity thresholding to segment individual nuclei (based on the most intense fluorescent channel between DNA or the fluorescent protein). The initial position of the bleached ROI was defined by its center coordinates and radius at the time of photobleaching. At subsequent time frames, the position of the bleached ROI was dynamically updated using a computer vision-based optical flow approach, implemented with pyramid Lucas-Kanade method (cv2.calcOpticalFlowPyrLK, OpenCV). Tracking was initialized using all pixel coordinates within the nuclear mask, and only correspondences with successful status values were retained. These good point correspondences were then used to compute an affine transformation matrix (cv2.estimateAffinePartial2D, OpenCV), which accounted for translational and rotational displacements of the nucleus. For quantification, mean fluorescence intensity was extracted for inside the bleached ROI (**S**) and outside the bleached ROI (**R**: reference) defined as the binary mask signal excluding a dilated version of the bleach ROI to avoid border effects. The recovery curve was calculated in each channel for all time points: 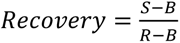 where **B** is the mean intensity of the background (86). Recovery curves for mKOet-H1, mEmerald-H3, and mEmerald-HP1α were temporally aligned to the first timepoint before photobleaching. DNA standard deviation / mean fluorescence intensity and roundness of nuclei were calculated at the first timepoint before photobleaching. Cells with poor nuclear masks or tracked bleached regions were excluded from further analysis. The recovery curves were fit to a model that describes diffusive behavior (87): 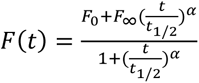. The fitted parameter 𝐹∞ and 𝐹_0_ were then used to calculate the mobile fraction (M_f_) as described in (86). were then used to calculate the mobile fraction (M_f_) as described in (86).

*<u>Quantification of H3 and HP1α dynamics:</u>* Time-lapse spinning disk confocal and DIC movies of NETosis-stimulated (Ionomycin calcium salt, 4 µM) dHL60 cells transfected with mEmerald-H3 or mEmerald-HP1α and stained with a DNA dye (SPY650-DNA) and a cell membrane dye (CellMask) were used to quantify the nuclear and cytoplasmic localization of these fluorescent proteins. For each time point, individual NETing cells were cropped and analyzed with a custom python script that applied an intensity thresholding to segment individual nuclei (based on the DNA channel) and individual whole cells (based on the CellMask channel). Cells with poor CellMask thresholds were segmented manually while cells with poor nuclear masks were excluded from further analysis. From these masks, we extracted features from each channel (H3, HP1α, DNA), such as standard deviation (SD), mean intensity, total intensity, area, with respect to time. We calculated cytosolic mean intensity as: (total cell intensity – nuclear total intensity)/ (total cell area – nuclear area). Each NETing cell was labeled for the timing when NETosis onset and nuclear rupture or end of the movie (whichever happens first). Single cell trajectories (SD, cytosolic or nuclear mean intensity) were temporally (x-axis) aligned so that they all start at NETosis onset and end 1 minute before nuclear rupture / end of the movie.

Amplitudes of single cell trajectories (SD, cytosolic or nuclear mean intensity) were subjected to a median filter and normalized relative to their values at NETosis onset enabling direct averaging and comparison across asynchronous cells. A Pearson correlation analysis (scipy.stats.pearsonr, SciPy) was performed between the cytosolic and nuclear mean intensities to assess divergence between these signals (coefficient value < 0). Pearson correlation coefficients were calculated using a 16-minute sliding window from NETosis onset to 1 minute before nuclear rupture. A peak-to-peak amplitude criterion was applied to ensure sufficient signal variability within each window.

*<u>Quantification of optical tweezer measurements</u>*: Time lapse brightfield and epifluorescence movies of NETosis stimulated dHL60 cells or U2OS cells stained with a plasma membrane dye (CellMask) and subjected to a tether pulling experiment were used (61). Trap forces (F) were calculated as the square root of the sum of x-force component squared and y-force component squared: 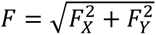 (88). A median filter was applied, and the steady state trap force was determined manually and averaged across 5-10 seconds. Tension was calculated using this formula: 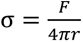 where r was determined by manually measuring the thickness of the tether, 2r, from the CellMask channel, using imageJ (derivation from ref (88)). The standard deviation of the nucleus (SD nucleus) was calculated by manually tracing the nucleus in the brightfield channel and using imageJ measure the standard deviation of the brightfield signal in the segmented area and normalizing it to the standard deviation of the background to minimize variability due to differences in illumination.*<u>Quantification of optical tweezer measurements</u>*: Time lapse brightfield and epifluorescence movies of NETosis stimulated dHL60 cells or U2OS cells stained with a plasma membrane dye (CellMask) and subjected to a tether pulling experiment were used (61). Trap forces (F) were calculated as the square root of the sum of x-force component squared and y-force component squared: 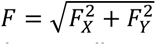 (88). A median filter was applied, and the steady state trap force was determined manually and averaged across 5-10 seconds. Tension was calculated using this formula: 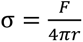 where r was determined by manually measuring the thickness of the tether, 2r, from the CellMask channel, using imageJ (derivation from ref (88)). The standard deviation of the nucleus (SD nucleus) was calculated by manually tracing the nucleus in the brightfield channel and using imageJ measure the standard deviation of the brightfield signal in the segmented area and normalizing it to the standard deviation of the background to minimize variability due to differences in illumination.

*<u>Quantification of volume measurements</u>*: 3D time lapse lattice light sheet movies of NETosis-stimulated dHL60 cells stained with a DNA dye (SPY650-DNA) and transfected with pmaxGFP were used to measure the evolution of cell volume during NETosis. We used the Imaris software to process the lattice light sheet CZI files and segment the cytosolic GFP signal in 3D with respect to time. Time dependent cell volume trajectories for individual NETing cells were extracted. The volume before microvesicles (MV) shedding (Bef. MV shedding) was defined as the average volume before ionomycin was added. The volume at the end of MV shedding and at the beginning of cell rounding (End of MV shedding) was defined as the average volume across 2-3 minutes centered at the onset of MV shedding. The volume before PM rupture (Bef. PM rupture) was defined as the average volume of the last 2-3 minutes before cell permeabilization or PM rupture as assessed by the loss of cytosolic GFP signal. The volume trajectories were temporally (x-axis) aligned between the following events: “Bef. MV shedding”, “End of MV shedding”, “Bef. PM rupture”. These trajectories were also normalized (y-axis) relative to “End of MV shedding” enabling direct averaging and comparison across cells.

**References Methods:** (Included in REFERENCES)

---

**Figure S1:**
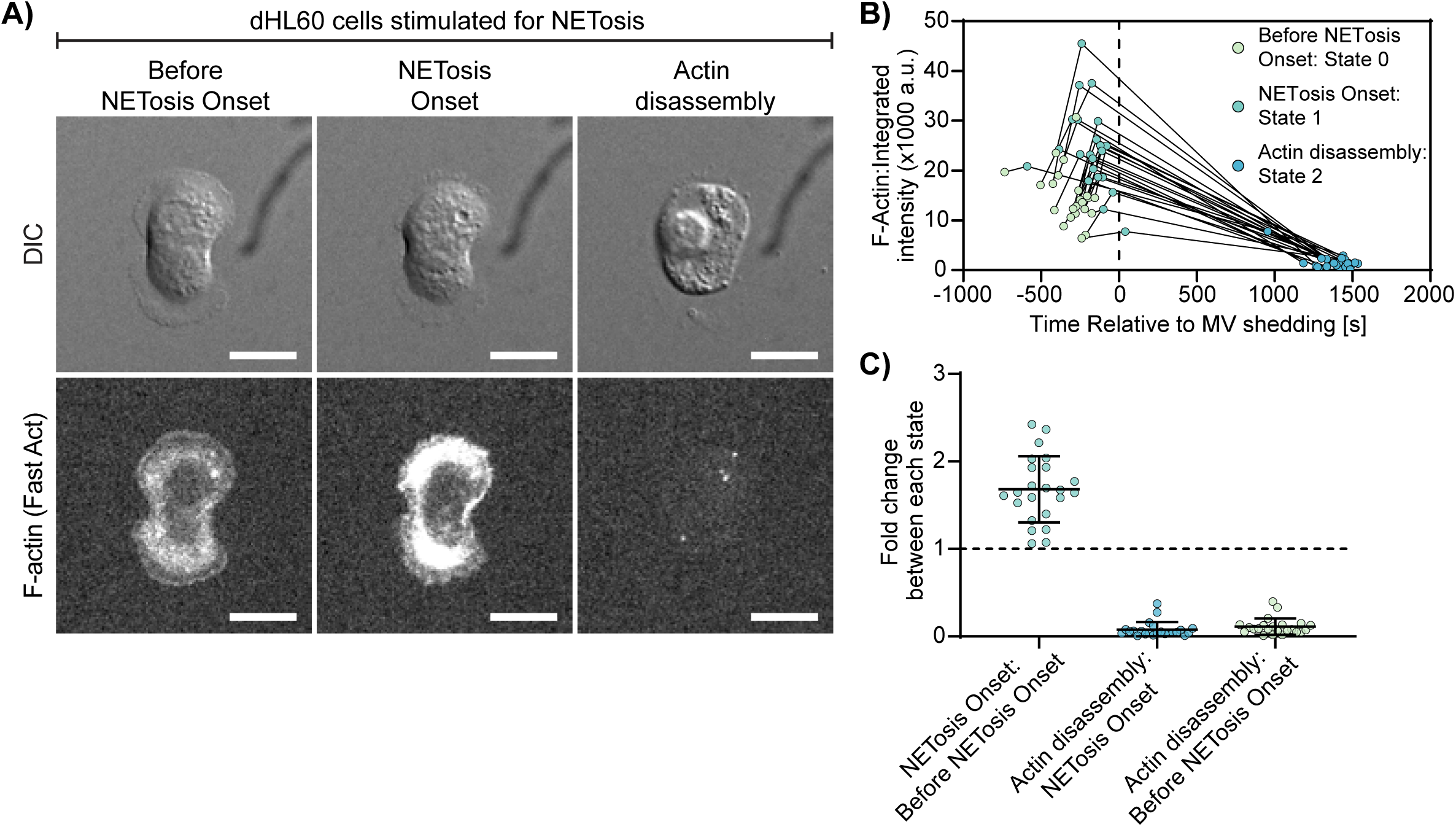
The actin cytoskeleton polymerizes before disassembling during early NETosis. dHL60 cells stained with FastAct (F-actin probe) were stimulated with ionomycin (4 μM) and imaged at the coverslip–cell interface (Z = 0 μm) by DIC and confocal microscopy at 5 second intervals for 30 min. **(A)** Representative montage of a dHL60 cell at key stages of NETosis : Before NETosis Onset, NETosis Onset (defined by microvesicle (MV) shedding and/or cell rounding), and Actin Disassembly. Top: DIC; Bottom: F-actin (FastAct) fluorescence. Brackets denote the presence of 4 μM ionomycin. Scale bar: 10 μm. **(B)** Single-cell trajectories of background-corrected integrated F-actin intensity over time, aligned relative to MV shedding (dashed line). States are indicated: before NETosis onset (green), NETosis onset (teal), and Actin Disassembly (blue). **(C)** Quantification of the fold-change of integrated F-actin intensity across states: NETosis Onset vs. Before NETosis Onset, Actin Disassembly vs. NETosis Onset, and Actin Disassembly vs. Before NETosis Onset. Each point represents a single cell; bars indicate mean ± SD. n = 23 cells. Actin disassembly is illustrated by the decrease in F-actin intensity in cell (**A**-bottom right image), and graphs **(B;** State 2) and (C; decreased fold change between “actin disassembly” and “NETosis onset” or “Before NETosis onset”). Note that the increased F-actin intensity in (A; NETosis Onset); (B; State 1); and (C; NETosis Onset : Before NETosis onset); suggests F-actin assembly.

**Figure S2:**
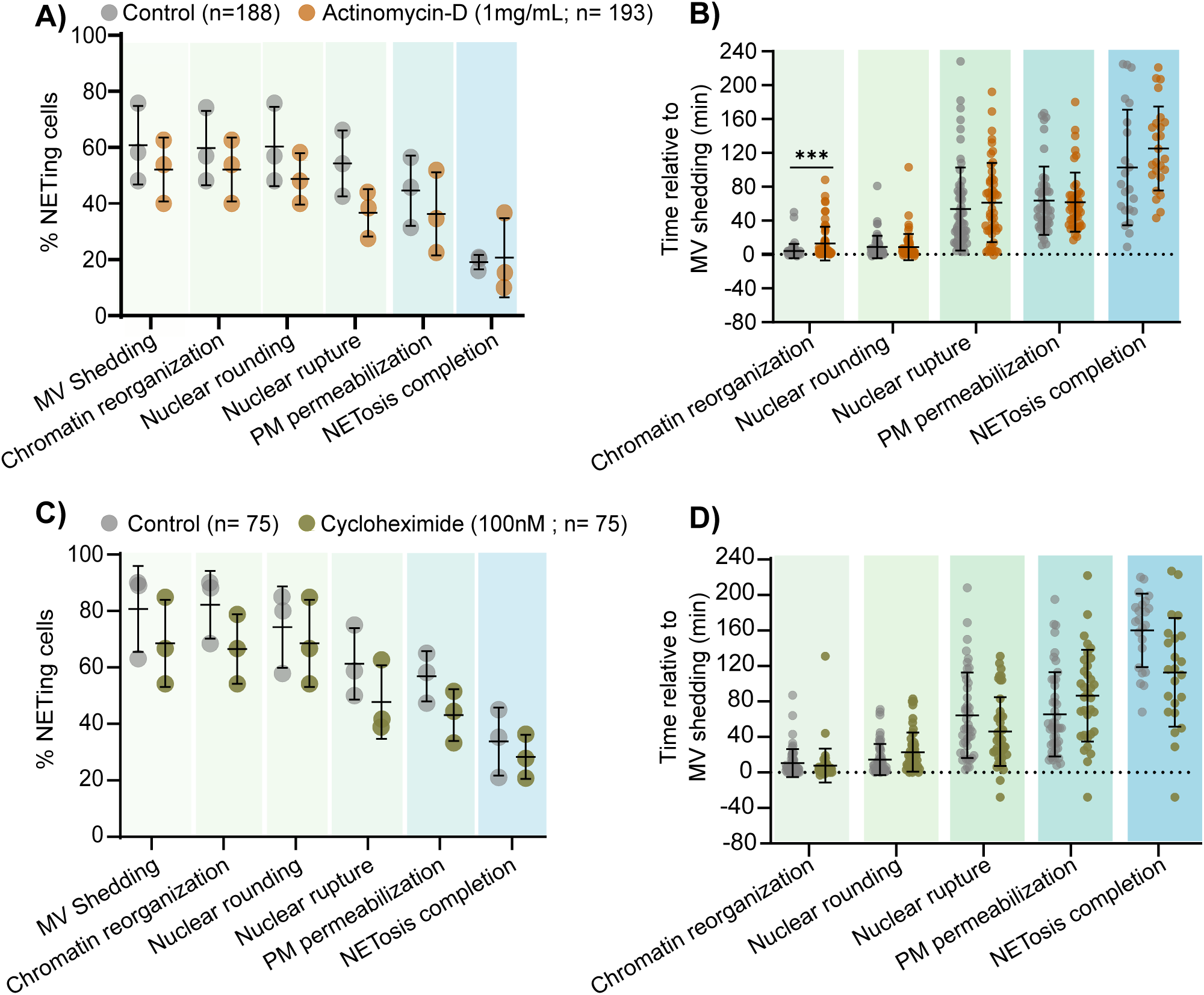
Active transcription and translation are not required for NETosis initiation and completion. dHL60 cells were treated with Actinomycin-D **(A-B)** or cycloheximide **(C-D)** for 1hr prior to being stained with SPY650-DNA and stimulated with ionomycin (4 µM) and imaged by DIC and confocal microscopy at coverslip-cell interface (Z= 0 μm) and in the cell center (Z =+3 μm) at 1 min intervals for 4h. Percentage of control and Actinomycin-D **(A)** or Cycloheximide **(C)** cells undergoing microvesicle (MV) shedding, chromatin reorganization, nuclear rounding, nuclear rupture, PM permeabilization, and NETosis completion after stimulation. n = total number of cells; each data point indicates percentage of cells from a biological replicate. **(B** and **D)** Timing of cellular events initiation relative to MV shedding. n = total number of cells, each data point indicates an individual cell. Grey datapoints represent untreated cells. Orange and green data points represent Actinomycin-D or Cycloheximide treated cells, respectively.

**Figure S3:**
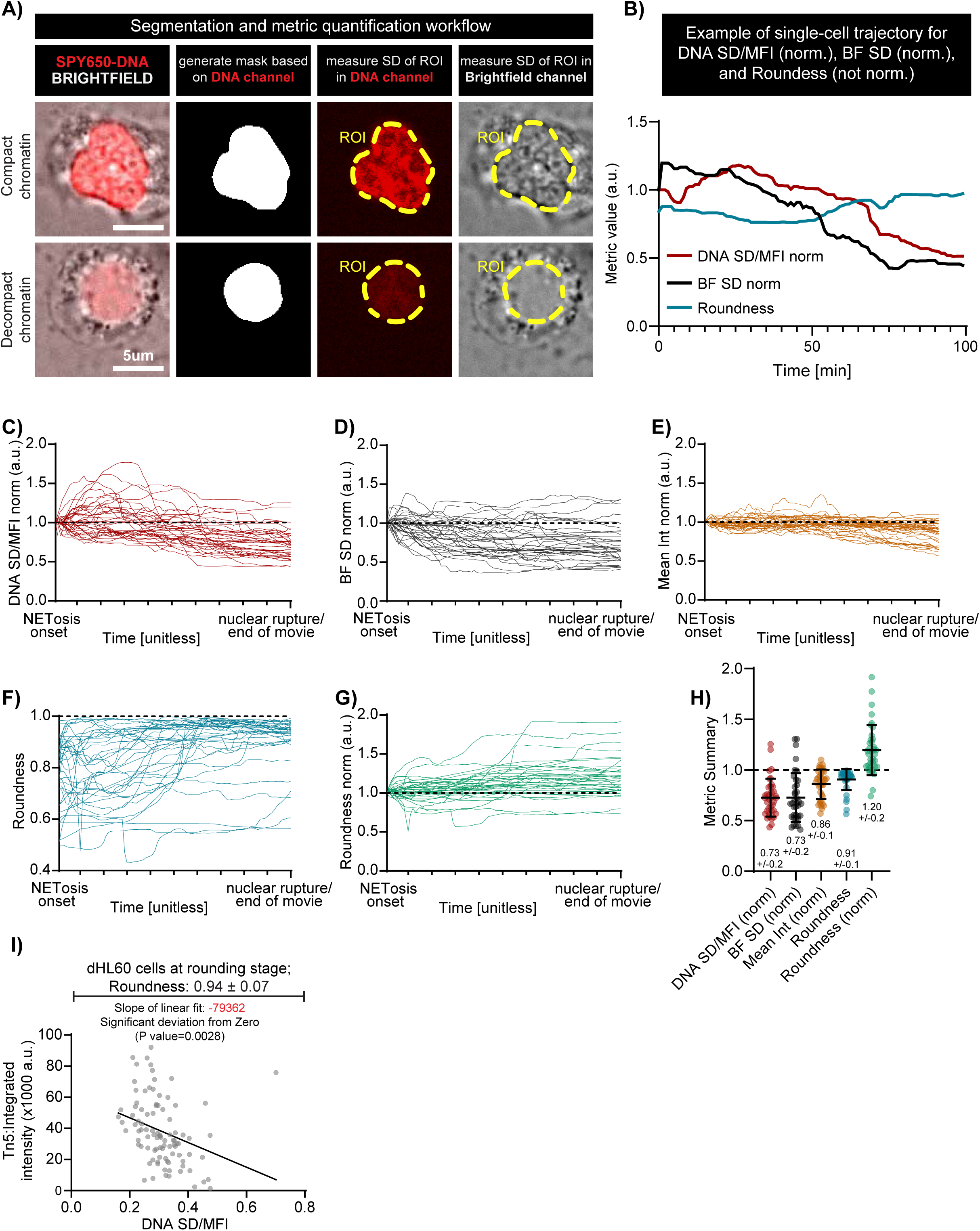
Nuclear signal homogenizes while roundness increases before nuclear rupture from NETosis onset. **(A)** Representative images and schematic of the single-cell segmentation workflow used to measure the standard deviation (SD) of SPY650-DNA and brightfield channels, the mean fluorescence intensity (MFI) in regions of interest (ROIs) as well as nuclear roundness between NETosis onset and nuclear rupture. Representative images of a NETosis-stimulated dHL60 cell with compact chromatin (Top) and decompact chromatin (Bottom). Scale bar: 5 μm. **(B)** Quantification of intensity and shape metrics of the cell in **(A)** tracked from NETosis onset until 1 min before nuclear rupture (100 min total). DNA SD/MFI (red) and Brightfield SD (black) were normalized to their initial value at NETosis onset, while Roundness (blue) was not normalized. **(C–G)** Single cell trajectories of intensity and shape metrics measured between NETosis onset and 1 min before nuclear rupture or end of the movie (whichever occurred first). Metrics include DNA SD/MFI **(C)**, Brightfield (BF) SD **(D)**, Mean Intensity (Int) **(E)**, and Roundness **(G)**, normalized (norm) to their initial value at NETosis onset, except Roundness **(F)**, which was not normalized. n = 37 cells from two biological replicates. **(H)** Quantification of the final values (1 min before nuclear rupture or end of movie, whichever occurred first) of DNA SD/MFI (red), Brightfield (BF) SD (black), mean intensity (int) (yellow), and Roundness (green), all normalized to their initial value at NETosis onset, except Roundness (blue). The dashed horizontal line at y = 1 is a visual guide for whether normalized metrics increased (>1), decreased (<1), or remained unchanged (≈1). n = 37 cells from 2 experimental replicates; bars indicate mean ± SD. **(I)** Quantification of Tn5 integrated intensity relative to DNA SD/MFI (stained with DAPI) at single cell resolution. Each dot represents a cell. Line in black represents a simple linear regression of chromatin homogenization (DNA SD/MFI) versus chromatin accessibility (Tn5 integrated intensity). Brackets denote the presence of 4 μM ionomycin. n = 95 fixed cells. Analysis performed using GraphPad Prism v10.

**Figure S4:**
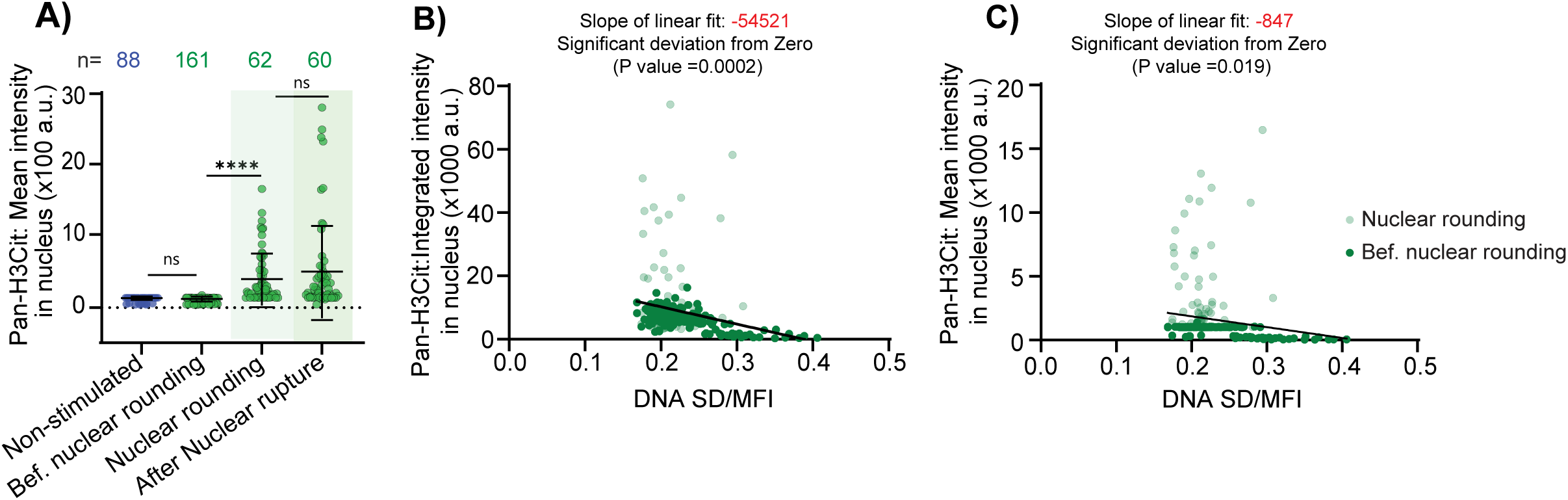
Histone H3 is citrullinated during early stages of NETosis. **(A)** Quantification of mean nuclear fluorescence intensity of Pan-H3Cit in non-stimulated and NETosis-stimulated dHL60 cells at different stages of NETosis as assessed by nuclear shape (roundness), nuclear envelope integrity (continuous or discontinuous LBR staining). n: total number of cells. Bars indicate mean ± SD. ns: not significant; **: p-value < 0.01 ****: p-value < 0.0001. Mean ± SD at the bottom. Statistical test; Mann-Whitney U test (two-tailed, non-parametric). **(B** and **C)** Quantification of integrated **(B)** and mean **(C)** Pan-H3cit fluorescence intensity relative to DNA SD/MFI at single cell resolution. Each dot represents a cell. Lines in black represent simple linear regression between chromatin homogenization (DNA SD/MFI) and Pan-H3Cit intensity. Slopes of linear regressions (indicated in red) are significantly different from zero indicating a significant negative correlation between the Pan-H3Cit intensity and chromatin homogenization. Analysis was performed using GraphPad Prism v10.

**Figure S5:**
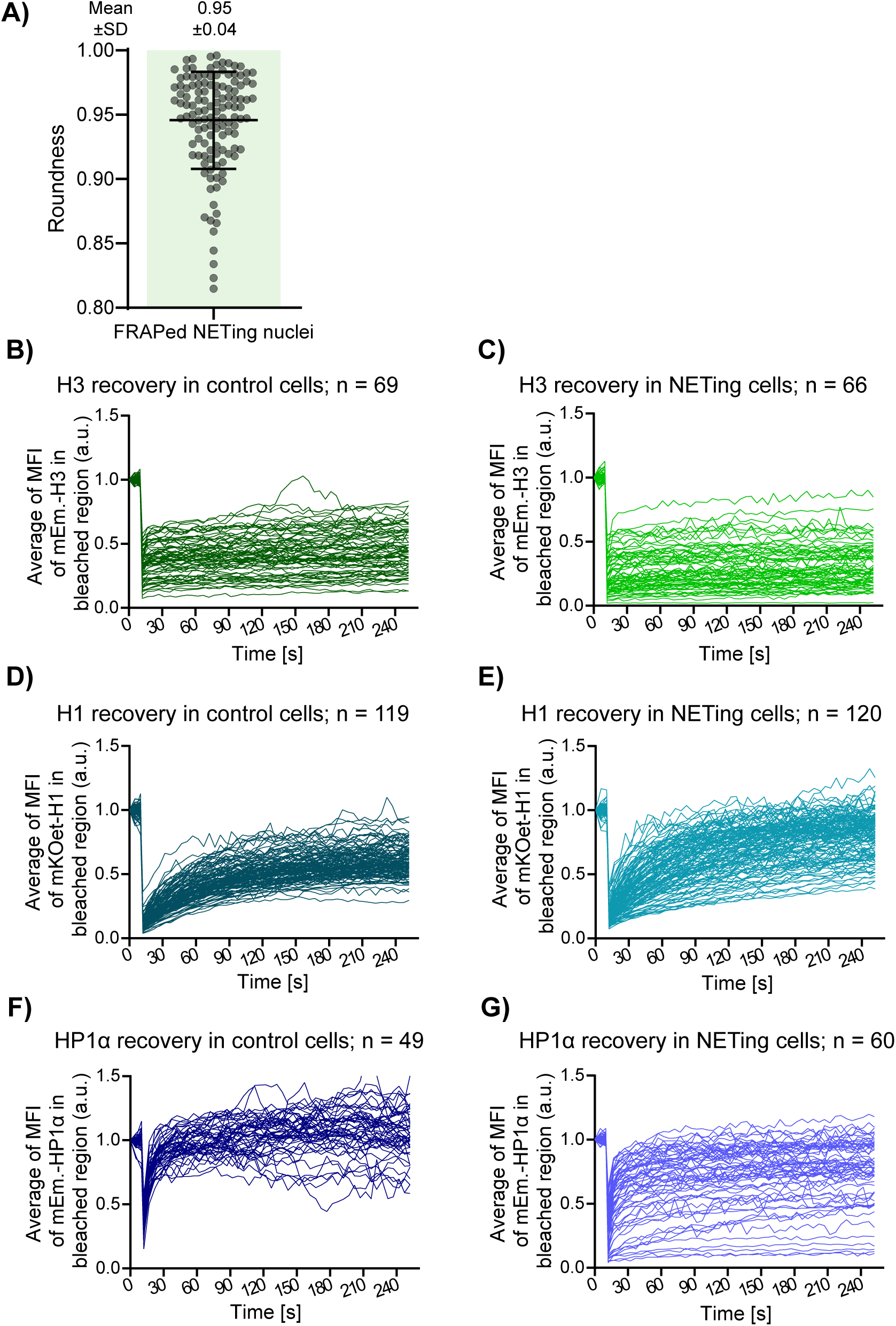
Dynamics of H3, H1, and HP1α recovery in the nucleus during NETosis. **(A)** Quantification of nuclear roundness in FRAPed NETing cells. Each dot represents a cell. Bars represent Mean ± SD and values are shown at the top. Green shaded rectangle indicates cells undergoing NETosis. FRAP recovery curves of individual dHL60 cells calculated as the mean fluorescence intensity (MFI) in the bleached nuclear region, corrected for background and normalized to the fluorescence intensity in the unbleached, reference region. Curves are shown for mEmerald-H3 in control **(B)** and NETing cells **(C)**, mKOet-H1 in control **(D)** and NETing cells **(E)**, and mEmerald-HP1α in control **(F)** and NETing cells **(G)**. n: number of cells.

**Figure S6:**
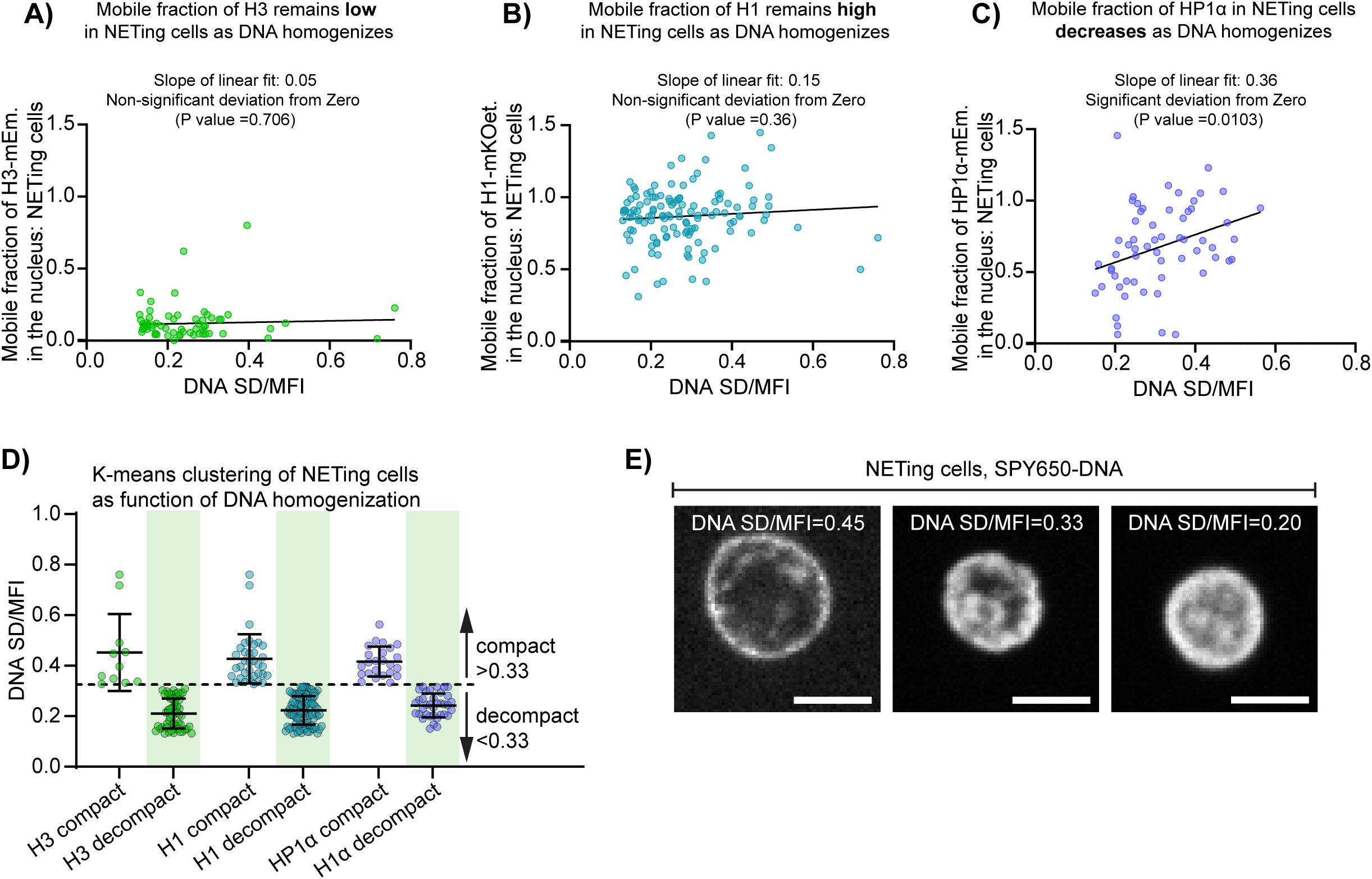
The mobile fraction of H1 and HP1α in the nucleus changes with chromatin decompaction (A-C) Correlation analysis between the mobile fractions of mEmerald-H3 **(A)**, mKOet-H1 **(B)** and mEmerald-HP1α **(C)** (shown in Figure 2K and L) and DNA SD/MFI (to assess chromatin homogenization). Lines in black represent simple linear regression between mobile fractions and DNA SD/MFI. Slopes of linear regression and their significant or non-significant deviation from zero are indicated above their respective graphs. **(D)** Quantification of DNA SD/MFI of FRAPed NETing cells stained with SPY650-DNA. DNA SD/MFI of all NETing cells (n = 252 cells) shown in Figure 2K and L were clustered in two groups using k-means clustering to identify that 0.33 is the value of DNA SD/MFI that best distinguishes cells with compact (DNA SD/MFI > 0.33) versus decompact (DNA SD/MFI > 0.33) chromatin. Green shaded rectangles indicate cells with decompacted chromain, defined as DNA SD/MFI < 0.33. Data are shown as Mean ± SD. **(E)** Representative images of SPY650-DNA from live dHL60 cells at different chromatin of chromatin compaction and corresponding DNA SD/MFI. Images correspond to the first time point before FRAP. Brackets denote the presence of 4 μM ionomycin. Scale bar: 5 μm. Regression analyses were performed in GraphPad Prism v10.

**Figure S7:**
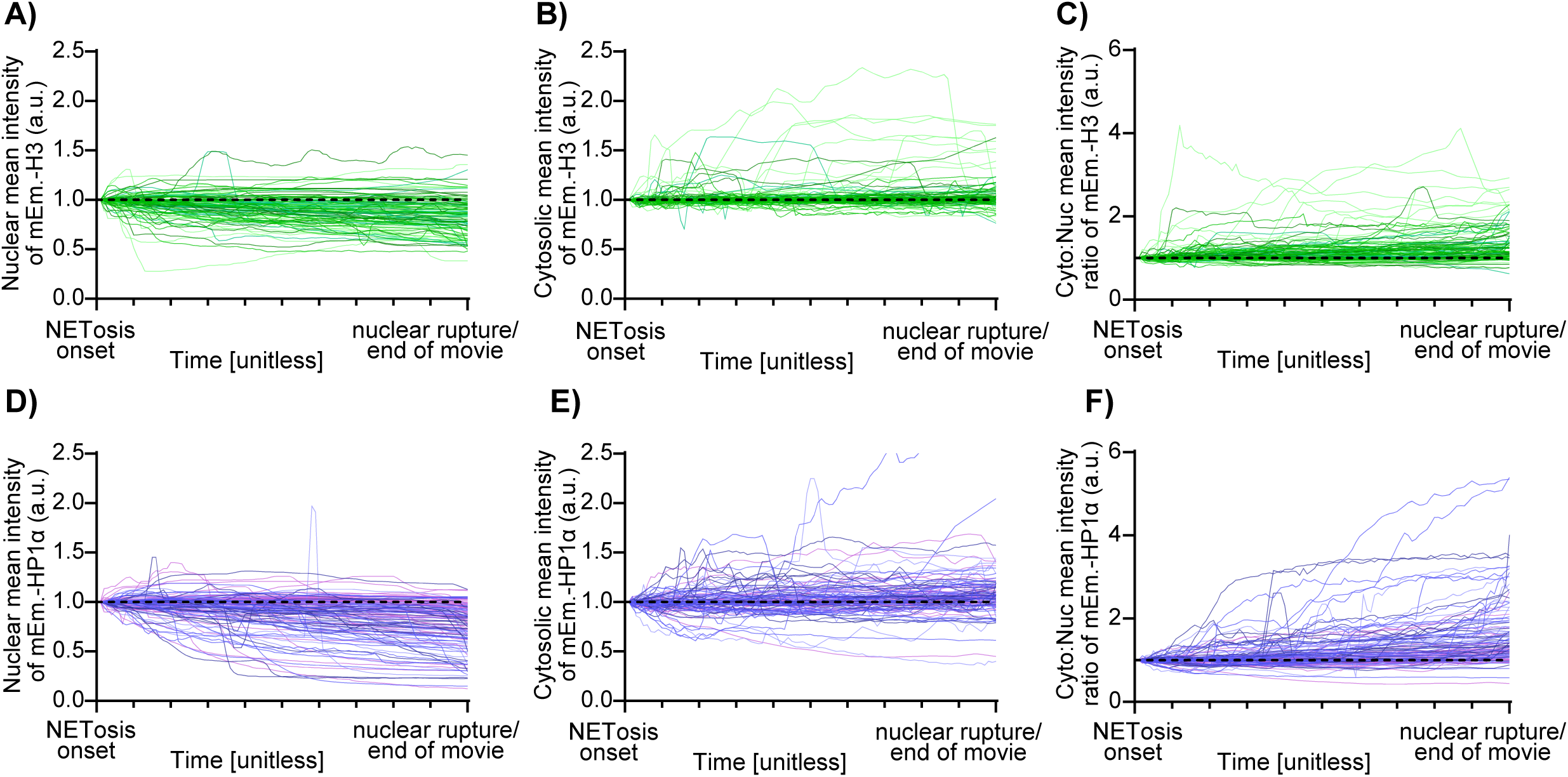
The dynamics of H3 and HP1α from MV shedding to nuclear rupture. Quantification of nuclear, cytosolic, and cytosolic:nuclear intensities of mEmerald-H3 **(A–C;** n = 115, 94, and 94 cells) or mEmerald-HP1α **(D–F;** n = 131, 111, and 111 cells) in dHL60 cells stimulated for NETosis. Intensity curves were temporally aligned to all start at NETosis onset (microvesicle shedding or onset of cell rounding) and end 1 min before nuclear rupture or end of the movie (whichever occurred first). Intensities were normalized to their initial value at NETosis onset. Different shades of green **(A-C**: mEmerald-H3) or purple **(D-F**: mEmerald-HP1α) represent four biological replicates.

**Figure S8:**
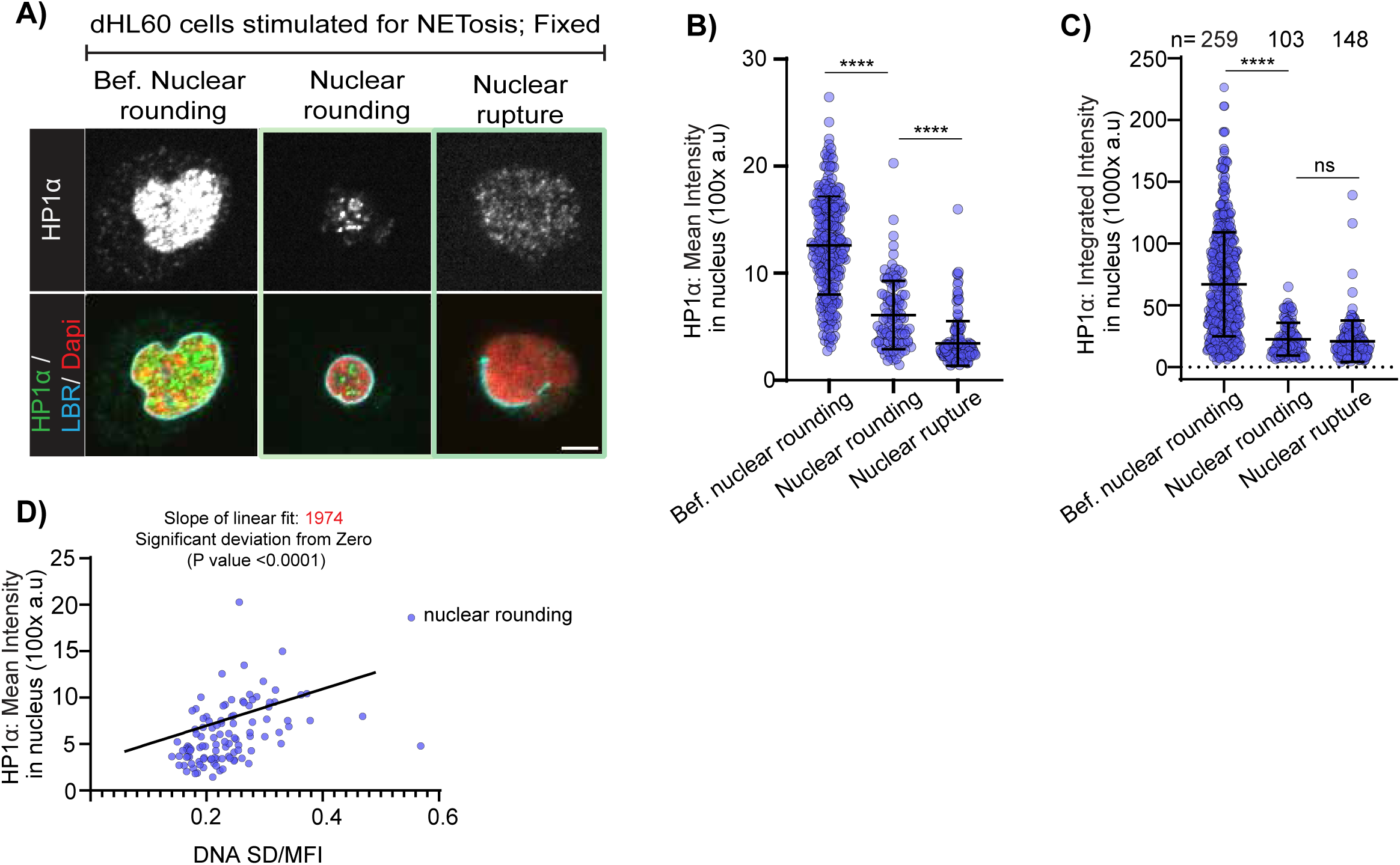
Endogenous nuclear HP1α decreases from the early stages of NETosis. **(A)** Representative immunostaining images of endogenous HP1α only (top) or overlayed with DAPI and lamin B receptor (LBR; Bottom) in dHL60 cells at different stages of NETosis, from 3D Z stack; step size 0.3 µm and range 9 µm. Brackets denote the presence of 4 μM ionomycin. **(B-C)** Quantification of integrated **(B)** and mean **(C)** fluorescence intensity of endogenous HP1α associated with DNA of cells at different stages of NETosis as assessed by nuclear shape (roundness), nuclear envelope integrity (continuous or discontinuous LBR staining). Each datapoint represents a cell. n: number of cells; Bars: Mean ± SD. **(D)** Correlation analysis between mean fluorescence intensity of endogenous HP1α in nucleus (data from **C**: nuclear rounding) and chromatin homogeneity (measured by DNA SD/MFI) in dHL60 cells at early NETosis (nuclear rounding). Line in black represents a linear regression with slope, indicated in red, is significantly different from zero indicating a significant correlation between HP1α intensity in the nucleus and chromatin homogenization. Each dot represents a single nucleus. ns: not significant; ****p-value < 0.0001. Statistical test; Mann-Whitney U test (two-tailed, non-parametric). Analysis was performed using GraphPad Prism v10.

**Figure S9:**
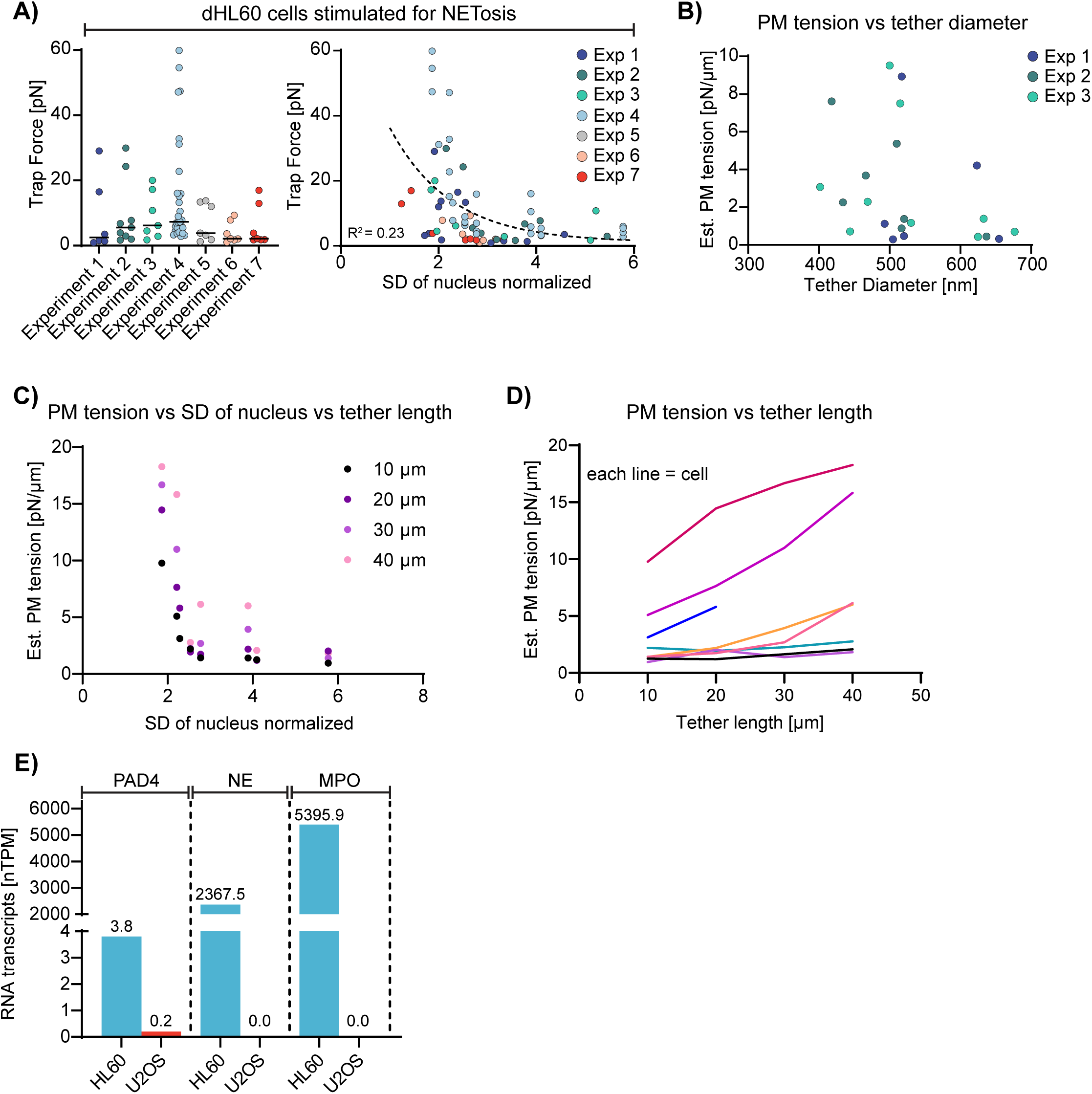
Plasma membrane tension increases during NETosis independently of the actin cytoskeleton and of membrane reservoir. **(A)** Quantification of trap force in NETosis-stimulated dHL60 cells, across multiple (7) experimental replicates, including the 4 replicates shown in Figure 4B and C. Data are shown as a column scatter plot (left) and as trap force plotted against the standard deviation (SD) of the nucleus in the brightfield channel, a metric of chromatin decompaction (right; Figure 1E). Dashed curve indicates the one-phase decay nonlinear regression fit of the data. Brackets denote the presence of 4 μM ionomycin. **(B)** Estimated (Est.) plasma membrane tension plotted as a function of tether diameter for replicates 1–3. **(C)** Quantification of estimated (Est.) plasma membrane tension as a function of chromatin decompaction (SD nucleus) at varying tether lengths (10, 20, 30, 40 μm), represented by different shades of purple. **(D)** Single-cell plots of estimated (Est.) plasma membrane tension versus tether length (same data as in **C)**, where each line represents an individual cell. **(E)** Graphical representation of data from the Human Protein Atlas, www.proteinatlas.org, (Uhlén M et al, 2015) showing the abundance of RNA transcript for PAD4, neutrophil elastase (NE), and MPO in HL60 and U2OS cells.

## Supplementary Movie Legends

Link to Google Drive folder

**Movie S1. Time-lapse movie of a representative dHL60 cell undergoing NETosis.**

Differentiated neutrophil-like (dHL60) cells were co-stained with SPY650-DNA (red, left panel) and CellMask (cyan, right panel). Differential interference contrast (DIC) and spinning-disk confocal images were acquired at 1-min intervals, 3 μm above the coverslip surface. Cells were stimulated with 4 μM ionomycin, and elapsed time (min) is shown relative to microvesicle shedding. Scale bar, 5 μm.

**Movie S2. Actin cytoskeleton disassembles during early NETosis**

Differentiated neutrophil-like (dHL60) cells were stained with F-actin dye (SPY555-FastAct). Differential interference contrast (DIC, left panel) and spinning-disk confocal (right, panel) images were acquired every 5 seconds for 30 minutes to monitor fast F-actin dynamics. Elapsed time is shown relative to microvesicle shedding. Scale bar, 10 μm

**Movie S3. Nuclear morphology and signal progressively homogenizes from NETosis onset to nuclear rupture**

Differentiated HL-60 (dHL60) cells were stimulated with 4 μM ionomycin and imaged using spinning-disk confocal microscopy. Three panels are shown: (left) merged brightfield and DNA channel (SPY650-DNA), (middle) DNA channel only, and (right) brightfield channel only. Images were acquired at 1-min intervals, 3 μm above the coverslip surface. Elapsed time (min) is indicated relative to NETosis onset. Scale bar, 5 μm.

**Movie S4. Representative FRAPed dHL60 cell expressing mEmerald-H3**

Differentiated neutrophil-like (dHL60) cells were transfected overnight (>10 hrs) with nucleosomal histone H3 fused to mEmerald (mEmerald-H3, gray, 1st column) and stained with SPY650-DNA. The 2nd column shows the merged signals (mEmerald-H3, cyan; DNA, red). Rows compare non-stimulated cells (top) and NETing cells (bottom) recovery after photobleaching a ∼3 μm^2^ nuclear region. Time-lapse spinning-disk confocal images were acquired at 5-s intervals. Elapsed time (sec) is shown relative to photobleaching. Scale bar, 5 μm.

**Movie S5. Representative FRAPed dHL60 cell expressing mKOet-H1**

Differentiated neutrophil-like (dHL60) cells were transfected overnight (>10 hrs) with linker histone H1 fused to mKOet (mKOet-H1, gray, 1st column) and stained with SPY650-DNA. The 2nd column shows the merged signals (mKOet-H1, cyan; DNA, red). Rows compare non- stimulated cells (top) and NETing cells (bottom) recovery after photobleaching a ∼3 μm^2^ nuclear region. Time-lapse spinning-disk confocal images were acquired at 5-s intervals. Elapsed time (sec) is shown relative to photobleaching. Scale bar, 5 μm.

**Movie S6. Representative FRAPed dHL60 cell expressing mEmerald-HP1α**

Differentiated neutrophil-like (dHL60) cells were transfected overnight (>10 hrs) with heterochromatin protein 1α fused to mEmerald (mEmerald-HP1α, gray, 1st column) and stained with SPY650-DNA. The 2nd column shows the merged signals (mEmerald-HP1α, cyan; DNA, red). Rows compare non-stimulated cells (top) and NETing cells (bottom) recovery after photobleaching a ∼3 μm^2^ nuclear region. Time-lapse spinning-disk confocal images were acquired at 5-s intervals. Elapsed time (sec) is shown relative to photobleaching. Scale bar, 5 μm.

**Movie S7. mEmerald-H3 exits the nucleus before nuclear rupture**

Differentiated neutrophil-like (dHL60) cells were transfected with nucleosomal histone H3 fused to mEmerald (mEmerald-H3, gray, 1st panel) and stained with SPY650-DNA (red, 2nd panel). The 3rd panel shows the merged mEmerald-H3 and DNA signals, and the 4th panel shows the merged DIC and DNA channels. Time-lapse DIC and spinning-disk confocal images were acquired at 1-min intervals, 3 μm above the coverslip surface. Elapsed time (min) is indicated relative to NETosis onset. Scale bar, 5 μm.

**Movie S8. mEmerald-HP1α exits the nucleus before nuclear rupture**

Differentiated neutrophil-like (dHL60) cells were transfected with heterochromatin protein 1α (HP1α) fused to mEmerald (mEmerald-HP1α, gray, 1st panel) and stained with SPY650-DNA (red, 2nd panel). The 3rd panel shows the merged mEmerald-HP1α and DNA signals, and the 4th panel shows the merged DIC and DNA channels. Time-lapse DIC and spinning-disk confocal images were acquired at 1-min intervals, 3 μm above the coverslip surface. Elapsed time (min) is indicated relative to NETosis onset. Scale bar, 5 μm.

**Movie S9. Classic tether pulling experiment on dHL60 cells stimulated for NETosis**

Differentiated neutrophil-like (dHL60) cells were stimulated with 4 μM ionomycin and a 2-μm, ConA coated carboxyl latex bead is optically trapped, gently touched to the plasma membrane and pulled away to form a tether. The steady state trap force and thickness of the tether is used to estimate the plasma membrane tension. Elapsed time, seconds. Scale bar, 5 μm.

**Movie S10. Trap force dynamics as a function of time during NETosis progression**

Differentiated neutrophil-like (dHL60) cells were stimulated with 4 μM ionomycin and a 2-μm, ConA coated carboxyl latex bead is optically trapped, gently touched to the plasma membrane and pulled away to form a tether. This bead is held stationary for ∼200 seconds to observe whether trap force remains constant throughout NETosis or changes during NETosis progression. Elapsed time, seconds. Scale bar, 5 μm.

**Movie S11. Lattice light sheet microscopy of dHL60 cell undergoing NETosis for volumetric measurements**

Differentiated neutrophil-like (dHL60) cells were transfected with pmaxGFP to express cytosolic GFP to visualize whole cell morphology during NETosis. Time-lapse light sheet images were acquired at 1-min intervals from before stimulation to right before plasma membrane rupture. Shown is an XYZ view of 3D render based on cytosolic GFP created with Imaris. Elapsed time, minutes. Scale bar, 10 μm.

